# A reference single-cell map of freshly dissociated human synovium in inflammatory arthritis with an optimized dissociation protocol for prospective synovial biopsy collection

**DOI:** 10.1101/2022.06.01.493823

**Authors:** Sam G. Edalat, Reto Gerber, Miranda Houtman, Tadeja Kuret, Nadja Ižanc, Raphael Micheroli, Kristina Burki, Blaž Burja, Chantal Pauli, Žiga Rotar, Matija Tomšič, Saša Čučnik, Oliver Distler, Caroline Ospelt, Snežna Sodin-Semrl, Mark D. Robinson, Mojca Frank Bertoncelj

## Abstract

Single-cell RNA-sequencing is advancing our understanding of synovial pathobiology in inflammatory arthritis. Here, we optimized the protocol for the dissociation of fresh synovial biopsies and created a reference single-cell map of fresh human synovium in inflammatory arthritis. We utilized the published method for dissociating cryopreserved synovium and optimized it for dissociating small fresh synovial biopsies. The optimized protocol enabled the isolation of a good yield of consistently highly viable cells, minimizing the dropout rate of prospectively collected biopsies. Our reference synovium map comprised over 100’000 unsorted single-cell profiles from 25 synovial tissues of patients with inflammatory arthritis. Synovial cells formed 11 lymphoid, 15 myeloid and 16 stromal cell clusters, including IFITM2+ synovial neutrophils. Using this reference map, we successfully annotated published synovial scRNA-seq datasets. Our dataset uncovered endothelial cell diversity and identified SOD2^high^SAA1+SAA2+ and SERPINE1+COL5A3+ fibroblast clusters, expressing genes linked to cartilage breakdown (SDC4) and extracellular matrix remodelling (LOXL2, TGFBI, TGFB1), respectively. We broadened the characterization of tissue resident FOLR2+COLEC12^high^ and LYVE1+SLC40A1+ macrophages, inferring their extracellular matrix sensing and iron recycling activities. Our research brings an efficient synovium dissociation protocol and a reference annotation resource of fresh human synovium, while expanding the knowledge about synovial cell diversity in inflammatory arthritis.

## Introduction

Chronic inflammatory arthritides, such as rheumatoid arthritis (RA) and the family of spondyloarthritides are clinically, radiologically, and molecularly diverse systemic autoimmune diseases that involve peripheral synovial joints and the axial skeleton^1, 2^. They affect an important proportion of the world population and are associated with significant morbidity, lifelong disability, extraarticular manifestations and shorter life expectancy^3–5^. Advances in therapy over the last decades have improved patient outcomes considerably and have pointed towards unique but also shared pathogenic pathways in different arthritis types^6, 7^. Despite these advances, the management of inflammatory arthritides faces critical challenges, including the development of precision medicine, targeting the stromal cell compartment, and discovering novel pro-resolution drugs^8–10^.

The synovial tissue, a delicate inner layer of the joint capsule, is the primary disease site in inflammatory arthritides, while spondyloarthritis types also affect the entheses^11, 12^. Given the central role of the synovium in inflammatory arthritides, understanding the identity and heterogeneity of its vital cellular components is quintessential in improving arthritis management and therapy^13, 14^. Single-cell RNA-sequencing (scRNA-seq) paralleled with the advancement in minimally invasive synovial biopsy^15^ has expanded our knowledge about synovial cell composition across arthritis types^16, 17^ and activity states^18^ (e.g., remission vs flare). Furthermore, combined with functional experiments, single-cell omics has been instrumental in establishing new concepts in inflammatory arthritis, including the break of synovial lining macrophage barrier^19^ and divergent specialisation of lining and sublining synovial fibroblasts into matrix-destructive and proinflammatory cell types^20, 21^.

Biobanking of viably frozen tissues simplifies retrospective sample selection and centralized tissue processing and scRNA-seq, decreasing the cost, minimizing batch effects, and facilitating multicenter studies^16, 22^. However, dissociating cryopreserved synovia may lead to inconsistent cell recovery and possible loss of sensitive, short-lived cell populations such as neutrophils, thereby introducing an inherent analysis bias. This bias could be overcome by isolation and analysis of cells from fresh prospectively collected synovial tissue samples. To date, most synovial scRNA-seq studies were carried out on the RA synovium, mainly using viably cryopreserved synovial tissues or pre-sorted synovial cell types^16, 23^ and the protocol for dissociating cryopreserved synovia has been optimized by Donlin and colleagues^22^. The largest scRNA-seq dataset from the fresh RA synovium, published by Stephenson and colleagues, comprises more than 20’000 unsorted scRNA-seq cell profiles from synovectomies of five RA patients^17^.

Our team investigates the pathobiology of human tissues affected by autoimmune diseases^24, 25^, with the first research phase focusing on protocol optimization and establishment of reference tissue scRNA-seq maps. We have recently published an optimized dissociation protocol and scRNA-seq map of the fresh and cultured human skin^24^. In the current paper, we present an optimized protocol for efficient dissociation of prospectively collected fresh synovial biopsies from patients with inflammatory arthritis. Furthermore, we created a single-cell reference dataset with annotation of synovial cells from fresh human synovium in inflammatory arthritis, serving as a research resource for the scientific community.

## Methods

### Patient recruitment

We enrolled 40 consecutive patients with different types of inflammatory arthritis, who were admitted to the Department of Rheumatology, University Hospital Zurich, Switzerland, for synovial biopsy and introduction/change of arthritis therapy. The ethics committee of Canton Zurich, Switzerland, approved the collection and analysis of patient data and synovial tissues. All patients signed the informed consent for participation in the study before inclusion. The arthritis diagnosis was set at the time of admission and was based on classification criteria for different arthritis types^2, 26, 27^ and clinical examination by a rheumatologist. UA diagnosis was set when arthritis did not meet the criteria for any defined arthritis type. In total, we collected 42 synovial tissues from 19 patients with RA, 6 patients with spondyloarthritis, 6 patients with psoriatic arthritis, and 9 patients with UA. Two RA patients donated biopsies from two different joints. These paired biopsies were processed as independent samples for downstream analyses. Synovial tissue samples originated from different joints, including knees (n=17), wrists (n=13), metacarpophalangeal joints (n=10) and sternoclavicular joints (n=2).

### Ultrasound-guided synovial tissue biopsy

We obtained synovial tissue by the ultrasound-guided synovial biopsy using the Quick core 18G × 6cm or Quick Core 16G 9cm biopsy needles and GE Logiq S7 ultrasound instrument^28^. Joints were evaluated for synovitis by ultrasound with GE Logiq S7 ultrasound instrument and multifrequency (5-13 MHz) linear array transducers, prior to biopsy. Ultrasound synovitis scoring (grayscale synovial hypertrophy, PD ultrasound parameters, semiquantitative grading 0-3) was performed in accordance with the EULAR-OMERACT guidelines^29^. Synovial tissue biopsy fragments were utilized for immunohistology, optimization of tissue dissociation protocol, scRNA-seq analysis or other downstream applications. Multiple synovial tissue fragments were collected from each donor. Specifically, for histological analysis, we scored a median of 7 (4-15) tissue fragments per joint, collected from different synovial tissue sites within each joint. For scRNA-seq analyses, we pooled a median of 11 (6-20) biopsy tissue fragments from the same joint to mitigate the known fragment-to-fragment variability and facilitate representative capture of the synovitis process in individual joints.

### Histological analysis

To identify the presence of synovial tissue in biopsied tissue fragments and to grade synovitis, biopsy fragments were formalin-fixed, paraffin-embedded, and sectioned (2μm) followed by haematoxylin-eosin, Giemsa and Elastica van Gieson staining of tissue sections. The identity of synovial tissue in biopsied fragments was confirmed based on the presence of a synovial lining layer and histologic features of the synovial sublining layer (fibrovascular or fatty tissue). For each tissue sample, grading of synovitis was performed based on the Krenn synovitis scoring (0-9) and its sub-scores evaluating synovial lining hyperplasia (0–3), stromal cell activation (0-3) and immune cell infiltration (0–3)^30^. For pathotype analysis^14^, slides were deparaffinized and automatically stained with specific antibodies against synovial cell markers (PECAM-1, DAKO JC/70a, 1:10; CD20, Ventana Roche L26, prediluted; CD3, Leica LN10, 1:500; CD68, DAKO A/S PG-M1, 1:50; CD138, DAKO A/S, MI15, 1:30 and CD15, DAKO A/S Carb-3, 1:100). We used the HRP-based DAB staining for antigen visualization and performed pre-treatments according to the manufacturer’s instructions (Ventana BenchMark). Histological analyses were conducted at the Department of Pathology, University Hospital Zurich, Switzerland by two experts, including a pathologist and a rheumatologist.

### Cell isolation from fresh synovial biopsies

Synovial biopsies used for protocol optimization and scRNA-seq were submerged in a sterile RPMI culture medium, transferred to the laboratory, and processed within 1-2 hours of collection. Forty-two fresh synovial biopsy samples were dissociated either with the protocol of Donlin et al.^22^ (protocol 1, n=26), or our optimized protocol (protocol 2, n=16) derived from Donlin et al.^22^ Briefly, synovial biopsies were minced and digested in a mixed enzymatic-mechanical dissociation including 100 μg/ml Liberase TL (Roche) and 100 μg/ml DNAse I (Roche); red blood cells were lysed (Red Blood Lysis buffer, Roche), and synovial cell pellets washed. For a detailed description of our protocol (protocol 2) and reagents see Supplementary Methods and Key Resource Table. Cell counts and viability were determined using a manual (Neubauer, Trypan Blue) or automated (Luna-FL, Dual Fluorescence Cell Counter, Logos BioSystems, Acridine Orange/PI, ThermoFisher) cell counting.

### Study workflow

All 42 synovial biopsies used in this study were processed fresh, and the presence of synovium was confirmed histologically for all included samples. Among the 42 dissociated synovial tissue samples, six samples were utilized for other downstream applications while eight samples were discarded because of the compromised cell viability/yield. Among 28 sequenced samples, we excluded three samples because of the limited quality of the scRNA-seq library/data. The final synovial scRNA-seq dataset included single cell profiles from 25 high-quality synovial cell suspensions obtained from patients with RA (n=15 biopsies), psoriatic arthritis (n=3 biopsies), spondyloarthritis (n=4 biopsies) and UA (n=3 biopsies), dissociated either with protocol 1 or protocol 2. In integrative protocol analysis, we integrated scRNA-seq data from RA and PsA synovial tissue samples, which were evenly distributed between the two protocols. All spondyloarthritis synovia were dissociated with protocol 1 and all UA synovia with protocol 2. Finally, we integrated the scRNA-seq data from 25 synovial tissue samples to create a reference single-cell map of the fresh human synovium in inflammatory arthritis.

### Single-cell RNA-seq library preparation and sequencing

Synovial single-cell libraries were prepared using the 3’ v3.0 and 3’ v3.1 protocols and reagents from 10X Genomics according to the manufacturer’s instructions targeting the encapsulation of up to 6000 cells. We created two independent paired libraries from samples 23, 26 and 28 as part of the quality control pipeline. The quality and quantity of cDNA and libraries were assessed with Agilent Bioanalyzer (Agilent Technologies). Diluted 10nM libraries were pooled in equimolar ratios, and the library pools were sequenced on the Illumina NovaSeq6000 sequencer (paired-end reads, R1=28, i7=8, R2=91, min. 40,000 -50,000 reads/cell) at Functional Genomics Center Zurich, the University of Zurich and ETH Zurich, Switzerland. A total of 6 separate sequencing runs on pooled scRNA-seq libraries, representing sequencing projects, were performed, as shown in **Table 1**. Two library pools (**Table 1**) combined synovial and skin scRNA-seq samples, the latter being part of the skin protocol optimization and reference skin mapping project^24^.

**Table 1.** Per sample percentage of reads discarded because of swapped barcodes together with the sequencing project identity number (ID) Removal of swapped barcodes was performed for library pools o24300 and o24793, which contained scRNA-seq libraries from skin and synovium. Skin samples are part of another project^24^ and are not listed.

| Sample | Proportion of reads discarded | Sequencing project ID |
| --- | --- | --- |
| Syn_Bio_023 | NA | o20549, o20550 |
| Syn_Bio_026 | NA | o20549, o20550, o23841 |
| Syn_Bio_028 | NA | o21475 |
| Syn_Bio_049 | NA | o21475 |
| Syn_Bio_050 | NA | o21475 |
| Syn_Bio_053 | NA | o23841 |
| Syn_Bio_054A | NA | o23841 |
| Syn_Bio_062 | NA | o23841 |
| Syn_Bio_064 | NA | o23841 |
| Syn_Bio_074 | NA | o23841 |
| Syn_Bio_077a | NA | o23841 |
| Syn_Bio_077b | NA | o23841 |
| Syn_Bio_078 | 8.32% | o24300 |
| Syn_Bio_079 | 3.00% | o24300 |
| Syn_Bio_081 | 3.36% | o24300 |
| Syn_Bio_083 | 3.09% | o24300 |
| Syn_Bio_084 | 3.07% | o24300 |
| Syn_Bio_087 | 5.54% | o24300 |
| Syn_Bio_091 | 1.11% | o24793 |
| Syn_Bio_092 | 0.99% | o24793 |
| Syn_Bio_093 | 1.20% | o24793 |
| Syn_Bio_096 | 1.14% | o24793 |
| Syn_Bio_098a | 1.20% | o24793 |
| Syn_Bio_098b | 1.40% | o24793 |
| Syn_Bio_099 | 1.03% | o24793 |

### Bioinformatics analysis of scRNA-seq data

Transcript count tables were generated from fastq files using cellranger count {cellranger} (version 6.0.0) and the reference Genome GRCh38.p13 (transcripts from GENCODE Release 32^31^ with genes coding for mitochondrial rRNA and tRNA, protein-coding RNA, rRNA and tRNA. Samples 23, 26 and 28 were each prepared as two independent libraries, sequenced in separate batches, and one of the sample 26 libraries was subsequently re-sequenced to reach the comparable sequencing depth across all libraries. Independent sample 23, 26 and 28 libraries were combined with cellranger aggr. Additionally, the re-sequenced library from sample 26 was combined with the corresponding sample data. Pooling a subset of synovial libraries with skin scRNA-seq^24^ libraries led to a limited barcode swapping^32^ with subsequent detection of a minor keratinocyte cluster in the synovial cell dataset. The swapped molecules and empty droplets were removed at the beginning of the analysis using the R package DropletUtils^33, 34^. The resulting filtered count matrix was then used for downstream analysis. No special enrichment for neutrophils was done^35^. The analysis was performed in R (version 4.0.3)^36^ using Bioconductor (version 3.12)^37^ packages. The preprocessing consisted of the following steps: (heterotypic) doublet detection and removal using scDblFinder^38^, cell filtering using scuttle^39^ and SampleQC^40^ and normalization (log-transformed normalized expression values) using scuttle. Highly variable genes were selected using scran^41^, dimensionality reduction was performed using PCA and intrinsicDimension^42^ (selection of PCA components to keep). The data from the two protocols were integrated using a mutual nearest neighbours approach as implemented in the R Bioconductor package batchelor. The batchelor package has been specifically created for the integration of datasets from different batches such as for example different experimental factors. Batches were removed by matching mutual nearest neighbours as described in [43]. Evaluation of batch integration was done using CellMixS^44^. Cell-type assignment was done in sequential steps. We identified the main synovial cell types first and then assigned cell subtypes within the main cell types individually. The first step of the main cell annotation was graph-based clustering (shared nearest neighbour graph and louvain clustering) with bluster^45^ followed by manual annotation (second step) with assistance from automatic cell type assignment using CellID^46^, cluster-specific/enriched genes and known marker genes. Cell subtype annotation followed the same steps as the main cell type annotation, starting from highly variable gene selection and using only the respective cell subset. Proportions of neutrophils were calculated by dividing the total number of neutrophils in a sample by the total number of cells in the respective sample. Marker genes were determined with pairwise Wilcoxon tests (using scran). We combined these marker genes with known cell markers and manually selected genes identified in the cluster comparison analysis based on the log-fold change expression (logFC>I1I) to create the heatmaps. Differential abundance analysis was conducted using edgeR^47^. Figures were generated with ggplot2^48^ and ComplexHeatmap^49^.

### Integration of scRNA-seq dataset with publicly available human synovial scRNA-seq data

Our in-house dataset was integrated with datasets from Wei K et al.^23^ (scRNA-seq profiles of sorted stromal cells from cryopreserved synovial tissues of six RA patients) and Stephenson W et al.^17^ (scRNA-seq data from non-sorted, fresh-dissociated synovial cells from 5 RA patients). The data from Wei K et al.^23^ was preprocessed as described above since fastq files were available. The data from Stephenson W. et al.^17^ was available as a filtered count matrix. The three data sets (in-house, Wei K. at al.^23^, Stephenson W et al.^17^) were integrated by keeping only the union of observed genes. The integrated data was processed in the same way as described above. Batch correction was performed using batchelor, which allowed us to calculate the corrected log-expression values that were then used for label transfer using SingleR^50^. As a reference, we used our analysed and annotated reference synovium map with single cell profiles from 25 fresh synovial tissue samples of patients with inflammatory arthritis.

### Data and code availability

The complete scRNA-seq dataset from 25 synovial tissue samples of patients with inflammatory arthritis is available in the ArrayExpress database (accession number E-MTAB-11791, https://www.ebi.ac.uk/arrayexpress/). The data from patients 28 and 50 have been previously deposited in the NCBI Gene Expression Omnibus as part of another publication^51^ and are accessible also through GEO SuperSeries accession number GSE181082. The code and instructions for the integrative analysis of scRNA-seq data generated by protocols 1 and 2 are available at https://retogerber.pages.uzh.ch/protocol_synovial. The code and instructions for the synovial single-cell reference map analysis are available at https://retogerber.pages.uzh.ch/synovialscrnaseq.

## Results

### Optimized dissociation protocol for fresh synovium minimizes the sample dropout rate

We utilized the protocol by Donlin and colleagues (protocol 1) as a starting reference for dissociating fresh synovial biopsies in our study. Donlin’s protocol was used across different single cell omics studies, however, it is primarily optimized for cryopreserved synovia^22^. Due to low cell yield and/or viability (median 1896 viable cells, range 0-9625), we lost eight of 26 prospectively collected tissue samples dissociated with protocol 1, resulting in a 31% sample drop-out rate. This data inferred that further optimizations of the original protocol might be needed for efficient dissociation of small fresh prospectively collected synovial biopsies. Specifically, we modified the original protocol to enhance the release of cells from synovial biopsies and minimize cell loss by refining the washing steps and volumes of reaction mixes (see **Suppl. Methods** for details). Krenn synovitis scores^30^, as a global measure of synovitis, did not significantly differ between the samples dissociated with protocol 1 and our modified protocol 2 (p=0.32, **Table 2**). We isolated a higher number of cells (p= 0.037, **Fig. 1a**) from samples dissociated with the protocol 2; both, the protocol 2 refinements and synovial tissue heterogeneity of consecutively recruited patients could contribute to this difference. Notably, synovial cell suspensions from protocol 2 samples demonstrated consistently high cell viability irrespective of isolated synovial cell yields (**Fig. 1b**). In contrast to protocol 1, no samples were lost from protocol 2 dissociations (p=0.039, Chi-square test with Yates’ correction). These results suggested that by slight modifications to the original protocol, we could decrease cell loss during sample processing and thus minimize the dropout rate of prospectively collected fresh synovial biopsies for scRNA-seq analyses.

**Figure 1.**
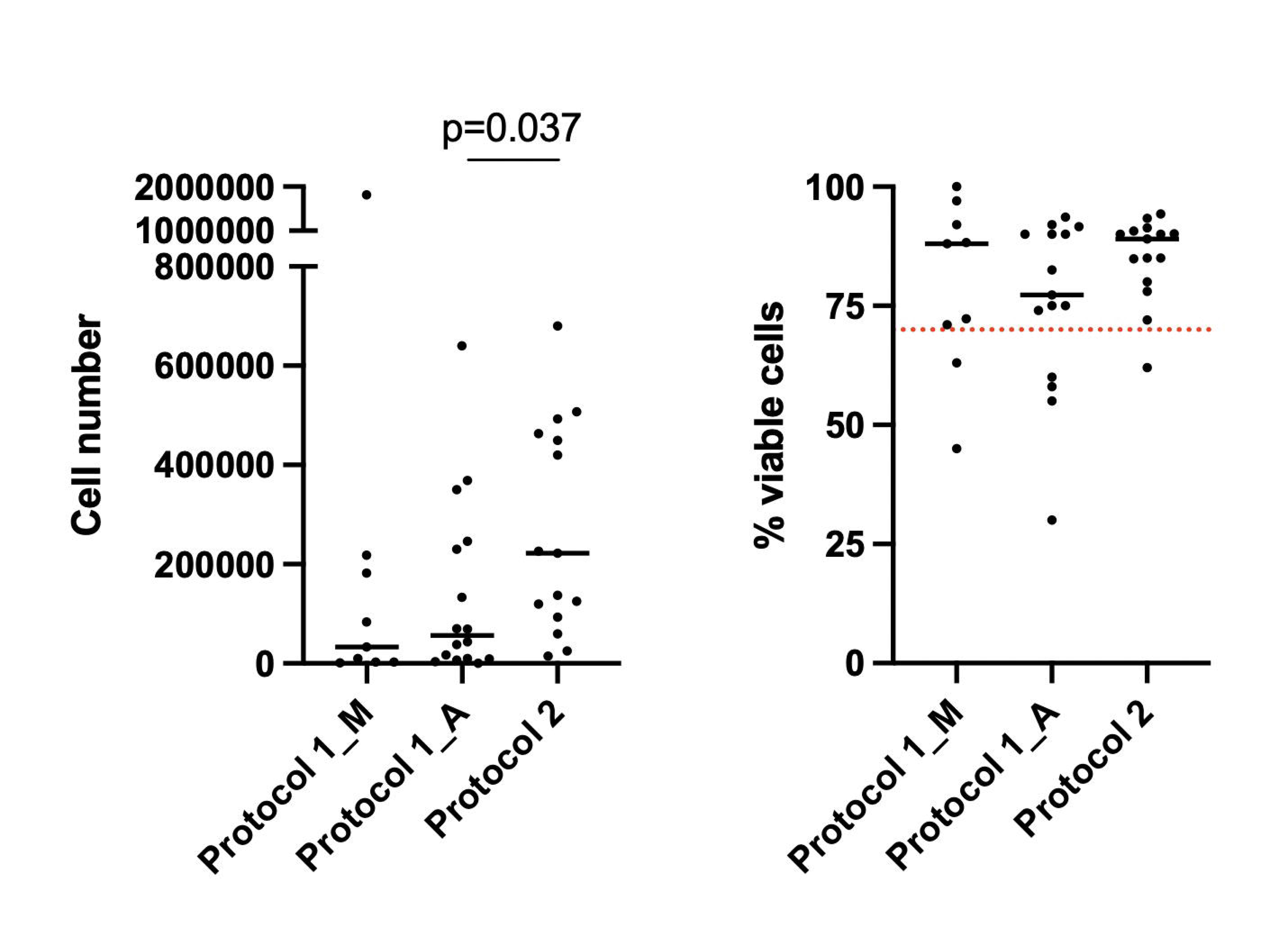
Number and viability of isolated synovial cells across synovial cell isolation protocols. Shown is the median. Protocol 1 – the original protocol of Donlin et al. Protocol 2 – the optimized protocol derived from Donlin et al. M: manual cell counting using the Neubauer chamber. A: automatic cell counting with the Luna-FL double fluorescence cell counter.Statistics: comparison of Luna counter outputs of cell yield and viability between the protocols, two-tailed Mann-Whitney test, *p=0.037.

**Table 2.** Histology characteristics and 10x Genomics chemistries for 18 synovial tissue samples from patients with inflammatory arthritis, included in the integrative analysis of scRNA-seq data between the original (protocol 1) and optimized (protocol 2) protocols. Numbers denote the biopsies processed. F: female, M: male, Pathotypes: DM: diffuse myeloid, LM: lymphoid myeloid, F: fibroid, pauci-immune, U: unclassified; *gender data not reported for 1 patient in protocol 2 cohort, **Krenn scoring in protocol 2 patient cohort based on 10 out of 12 synovial tissues. Statistics: Graph Pad Prism software, ***two-sided Fisher’s exact test, ****two-sided Mann-Whitney t-test, p< 0.05 accepted as a statistically significant difference.

|  | Protocol 1 (n=6) | Protocol 2 (n=12) | p value |
| --- | --- | --- | --- |
| Gender* |  |  | 0.24*** |
| Female | 6 | 7 |  |
| Male | 0 | 4 |  |
| Age* |  |  | 0.14**** |
| Median (range) | 71.5 (39 – 81) | 56 (20–72) |  |
| Krenn synovitis score** |  |  | 0.32**** |
| Median (range) | 5 (2-9) | 4 (2–6) |  |
| Pathotype |  |  | NA |
| Diffuse myeloid | 2 | 3 |  |
| Lympho-myeloid | 2 | 6 |  |
| Fibroid | 1 | 3 |  |
| Unclassified | 1 | 0 |  |
| 10x Genomics Chemistry<br>3' v3.0 | 5 | 0 | 0.0007**** |
| 3' v3.1 | 1 | 12 |  |

Given minute amounts of the biopsied material, we could not process paired tissue fragments from the same donor by both protocols, however unequivocal protocol-to-protocol comparison was not the objective of our study. Rather, we aimed to map fresh human synovium in inflammatory arthritis at a single-cell resolution and original protocol refinements facilitated our study goals.

### The optimized protocol enables detailed mapping of the human synovium

We created scRNA-seq data from unsorted synovial cells from 25 highly viable single-cell suspensions (median cell viability 90%, 24/25 samples with cell viability ≥ 70%, the threshold, below which viable cell sorting is highly recommended for scRNA-seq studies^52^). To understand whether our modifications to the original protocol led to differences in the synovial scRNA-seq data quality or cell type enrichment, we integrated scRNA-seq data generated by protocols 1 and 2. The integrative protocol analysis included 18 of 25 synovial tissue samples, thereby avoiding the diagnosis bias between the two protocols (see Methods for details). Patient demographic data, histology characteristics of the 18 synovial samples and utilized 10x Genomics chemistries are shown in **Table 2**.

Using highly-viable cell suspensions from protocols 1 and 2, we generated very good yields of scRNA-seq data that passes standard scRNA-seq QC filters (**Suppl. Fig. 1**). Specifically, we filtered out cells with a low total number of counts, a low total number of detected genes and a high percentage of mitochondrial counts (**Suppl. Fig. 1a**). The median numbers of cells kept per sample and detected genes per sample were 4467.5 and 14921, respectively (**Suppl. Fig. 1b**). The distribution of the total number of counts per cell (**Suppl. Fig. 1c**) and the total number of detected genes per cell (**Suppl. Fig. 1d**) was similar between samples dissociated with protocols 1 and 2.

We generated 76’902 high-quality scRNA-seq profiles from 18 synovial tissue samples that formed 11 principal synovial cell clusters comprising CD45+ lymphoid, CD45+ myeloid and CD45-stromal synovial cell subsets (**Fig. 2a**). The main synovial cell populations were heterogeneously distributed across patients’ synovia (**Fig. 2b)**; their abundances, however, did not differ between protocols 1 and 2 (**Fig. 2c**). The heatmap with top gene markers, specific for the 11 principal synovial cell populations, is shown in **Fig. 3**. Specifically, we identified synovial CD3E+ T cells, CD79A+ CD79B+ B cells, CD79A+ XBP1+ B cells/plasmablasts (**Suppl Fig. 2a**), CD14+ macrophages, LILRA4+ GZMB+ ITM2C+ plasmacytoid dendritic cells (DCs) (**Suppl Fig. 2b**), CD1C+ FCER1A+ DCs, CLEC9A+ DCs, CPA3+ mast cells, IFITM2^high^ neutrophils (**Suppl Fig. 2c**), VWF+ endothelial cells (ECs), ACTA2+TAGLN+ pericytes/mural cells and COL1A1+ synovial fibroblasts (**Suppl. Figs. 2d**). CD1C+ FCER1A+ and CLEC9A+ DCs co-clustered with synovial macrophages (**Suppl. Fig. 2b, c**), while CD3^neg^ NKG7+ NK cells co-clustered with T cells (**Suppl. Fig. 3a**).

**Figure 2.**
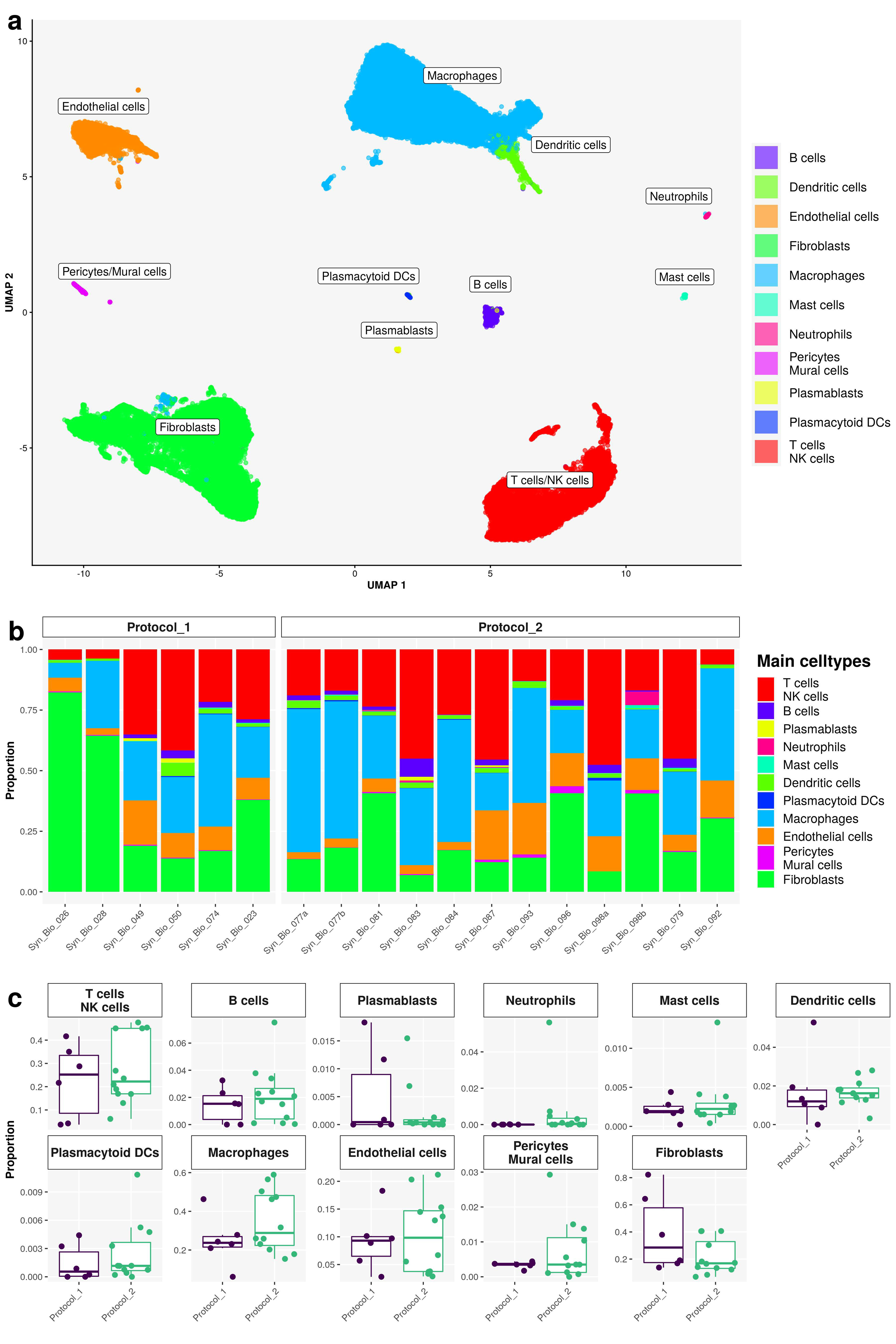
Integrative protocol analysis of scRNA-seq data. **a)** UMAP of the integrated scRNA-seq dataset with main synovial cell populations coloured by main cell type. ScRNA-seq data generated from fresh human synovia dissociated either with the original protocol of Donlin et al. (protocol 1, n=6) or the optimized protocol (protocol 2, n=12). **b)** Bar plots of relative abundances of main cell types per sample per protocol. **c)** The proportion of cell types per protocol 1 versus protocol 2; neutrophils were detected primarily in protocol 2 scRNA-seq data with a prominent enrichment in a subset of samples.

**Figure 3.**
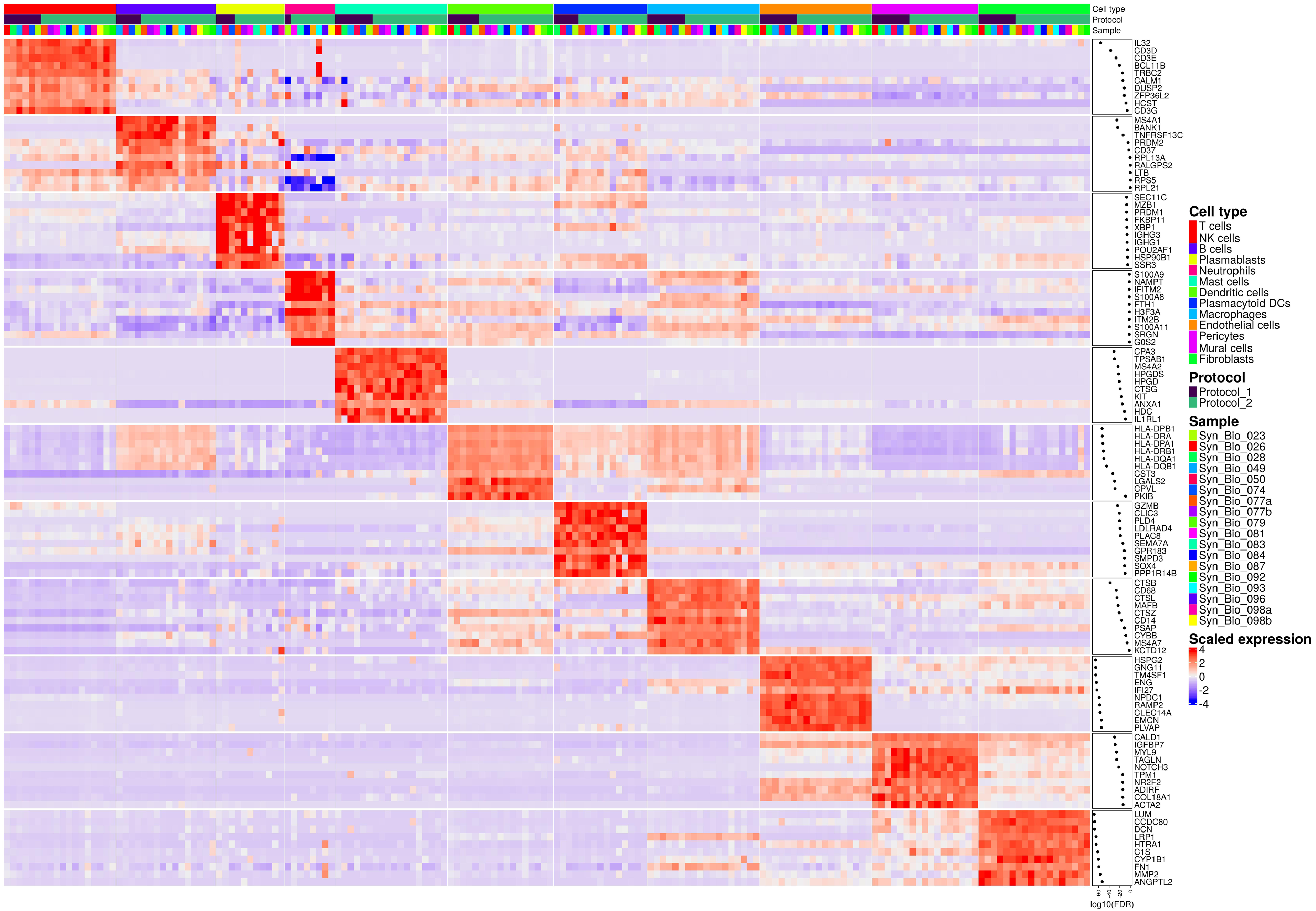
Top marker genes of the main synovial cell types identified in the integrative protocol scRNA-seq analysis. ScRNA-seq data generated from fresh human synovia dissociated either with the original protocol of Donlin et al. (protocol 1, n=6) or the optimized protocol (protocol 2, n=12). Shown is the heatmap with the top 10 cell type marker genes. Expressions are aggregated by sample and cell type.

Synovial T cells, macrophages, ECs, and synovial fibroblasts represented 4-48%, 6-59%, 3-21% and 7-82% of all synovial cells across the patient samples, constituting the most abundant synovial cells populations (**Fig. 2**). Looking closer at these cell populations, we identified CD4+ T cells, CD8A+ NKG7+ T cells (**Suppl. Fig. 3a**), tissue-resident synovial macrophages (expressing MERTK, TREM2 CD206/MRC1 genes, **Suppl. Fig. 3b**), infiltrating synovial macrophages (expressing IL1B, CD48 genes, Suppl. Fig. 3b), LYVE1^high^CCL21^high^ lymphatic ECs, vWF+ vascular ECs (**Suppl. Fig. 3c**), PRG4^high^THY1^low^ lining synovial fibroblasts, and THY1+ PRG4^low^ sublining synovial fibroblasts expressing GGT5, CXCL12 or DKK3 genes (**Suppl. Fig. 3d**).

Neutrophils were detected in 17% (1/6) of protocol one scRNA-seq samples and 50% (6/12) of protocol 2 samples (**Suppl. Table 1**), with stronger enrichment in a subset of protocol 2 samples (**Fig. 2c**). Top neutrophil genes are shown in **Suppl. Fig. 4**. Analyzing tissue histologically, we demonstrated neutrophil presence in 66% of protocol 1 samples and 91% of protocol 2 samples (**Suppl. Table 1**). In general, neutrophils were rather scarce and commonly focally distributed within or between biopsy fragments (**Suppl. Table 1**). Histology and scRNA-seq analyses matched in 50% when detecting synovial neutrophil presence/absence (**Suppl. Table 1**).

Overall, we could map all major and minor synovial cell populations with both protocols, inferring that our optimized protocol kept the comprehensiveness of the original protocol in detecting synovial cell diversity.

### Generation of a single-cell reference map of fresh human synovium in inflammatory arthritis

In the next step, we integrated the scRNA-seq data from 25 synovial tissue samples of patients with inflammatory arthritis (see Methods for details), generated either with protocol 1 or 2, to build a single-cell reference map of freshly dissociated human synovium in inflammatory arthritis. **Table 3** shows patient demographic data, histology characteristics and 10x Genomics chemistries for 25 synovial samples included in the human synovium reference map analysis. The QC analysis of the scRNA-seq data from the 25 samples is shown in **Suppl. Fig. 5**. After filtering the cells with a low total number of counts, a low total number of detected genes and a high percentage of mitochondrial counts (**Suppl. Fig. 5a**), we kept 102’758 single-cell profiles (**Suppl. Fig. 5b**). The distribution of the total number of counts per cell and the total number of detected genes per cell across the 25 synovial tissue samples is shown on **Suppl. Figs. 5c, d**. A single-cell map of fresh synovium comprised eleven principal synovial cell clusters (**Suppl. Fig. 6a**), which replicated the lymphoid, myeloid, and stromal cell types identified in the integrative protocol analysis (**Fig. 2a**). The variable distribution of these principal cell types across patient synovia is presented in **Suppl. Fig. 6b, c**. The top 10 marker genes defining these principal lymphoid, myeloid and stromal synovial cell populations are shown in **Suppl. Fig. 7**. Lymphocytes T, macrophages, synovial fibroblasts and ECs represented the most abundant cell types (**Suppl. Figs. 6**). We further subclustered these populations to understand their subset diversity and distribution in the synovium from patients with inflammatory arthritis.

**Table 3.**
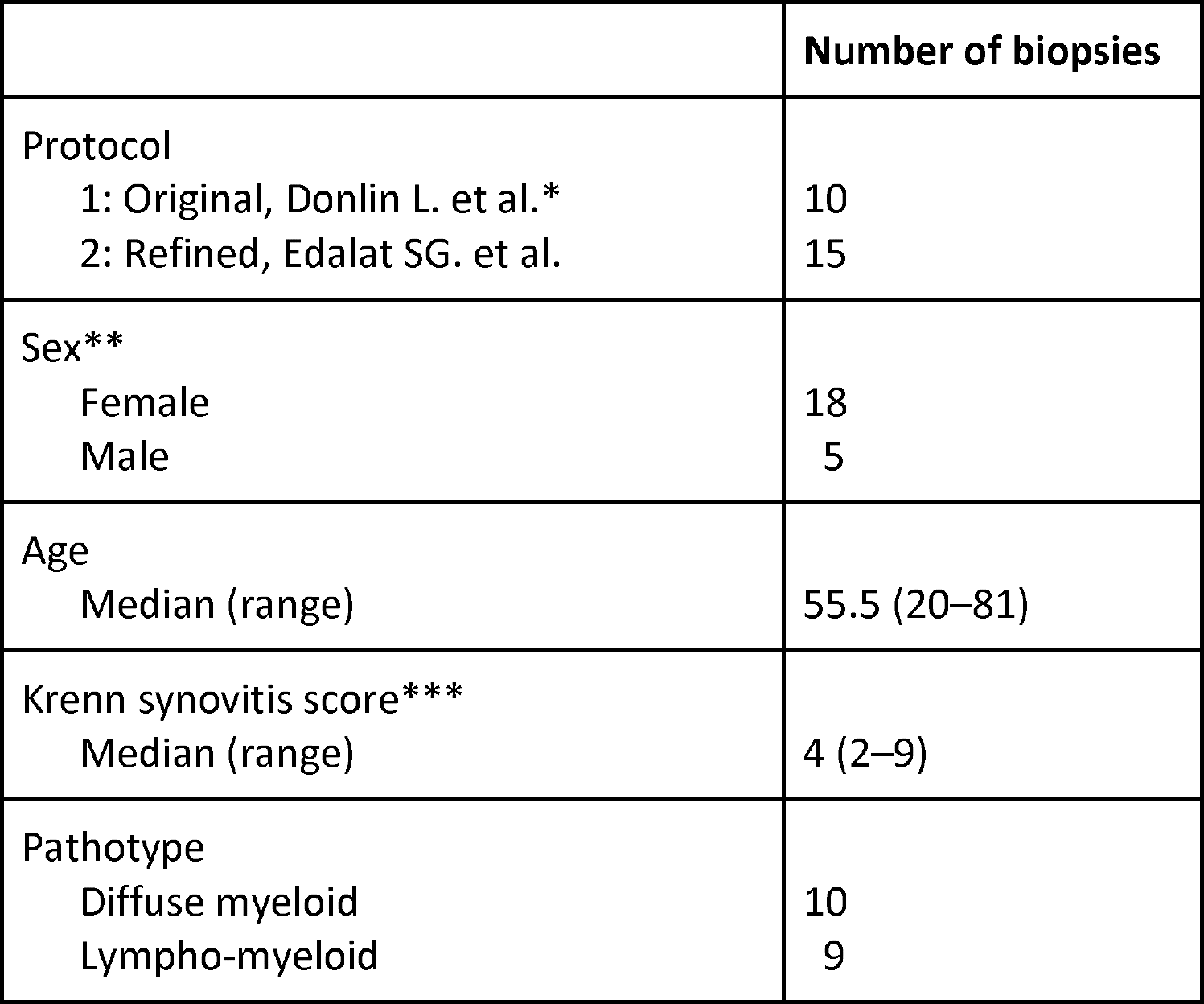
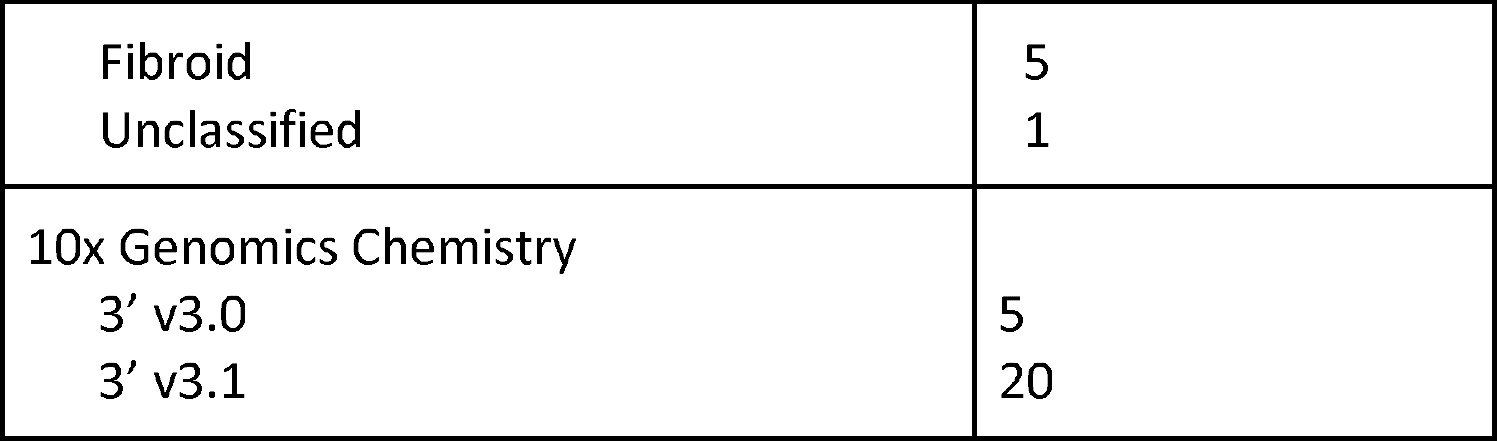
Histology characteristics and 10x Genomics chemistries for 25 synovial tissue samples from patients with inflammatory arthritis, included in the generation of the single-cell reference map of the fresh human synovium. Numbers denote the biopsies processed. F: female, M: male, DM: diffuse myeloid, LM: lymphoid myeloid, F: fibroid, pauci-immune, U: unclassified. *Donlin L. et al. Arthritis Res Ther 2019. **Gender data not reported for 2 patients in protocol 2 cohort, ***Krenn scoring based on 21 out of 25 synovial tissues.

|  | Number of biopsies |
| --- | --- |
| Protocol |  |
| 1: Original, Donlin L. et al.* | 10 |
| 2: Refined, Edalat SG. et al. | 15 |
| Sex** |  |
| Female | 18 |
| Male | 5 |
| Age |  |
| Median (range) | 55.5 (20–81) |
| Krenn synovitis score*** |  |
| Median (range) | 4 (2–9) |
| Pathotype |  |
| Diffuse myeloid | 10 |
| Lympho-myeloid | 9 |
| Fibroid | 5 |
| Unclassified | 1 |
| 10x Genomics Chemistry |  |
| 3' v3.0 | 5 |
| 3' v3.1 | 20 |

### Subcluster analysis uncovers the diversity of T cells in freshly dissociated human synovium

We profiled 23’169 cells comprising six CD3+ T cell clusters (1, 4-8), one cluster of CD3 ^neg^ NK cells (cluster 3), one cluster of proliferating TOP2A+ CENPF+ T cells and NK cells (cluster 2, **Suppl. Fig. 8a**) and a small cluster of CD3 ^neg^NKG7^neg^KLRB1+IL7R+ cells, suggestive of innate lymphoid cells (cluster 9) (**Fig. 4a**). The percent distribution of T cell, NK cell and innate lymphoid cell clusters varied considerably across patient’s synovia (**Fig. 4b, c**). The selected key marker and cluster-enriched genes of T cells, NK cells and innate lymphoid cells are presented in **Fig. 4d** and **Suppl. Fig. 8b**. Specifically, the CD4+ TIGIT ^low/neg^ T cell clusters (1, 5) differed in the expression of CCR7, LEF1, and SELL transcripts (**Fig. 4d, Suppl. Fig. 8c**). Meanwhile, CD4+ TIGIT+ CTLA4+ cluster 4 contained FOXP3+ and CXCL13+ PDCD1+ cells (**Figs. 4d, Suppl. Fig. 8c**), which could represent regulatory and peripheral/follicular CD4+ helper T cell phenotypes^16^. The CD8+ CCL5+ T lymphocytes comprised NKG7^med^ GZMK+ cells (cluster 6), cytotoxic NKG7^high^ GNLY^high^ GZMK^neg^ (cluster 8) and NKG7^high^ GNLY^neg^ GZMK^high^ (cluster 7) cells (**Figs. 4a, c, Suppl. Fig. 8d**). CD8+ T cell subsets varied in the expression of GZMB and GZMH transcripts (**Fig. 4d, Suppl. Fig. 8d**). The NKG7 ^high^ GNLY^high^ NK cells (cluster 3) were GZMB ^high^ and expressed abundantly Killer cell Lectin like Receptor (KLRC1, KLRD1, KLRF1) transcripts (**Fig. 4d**).

**Figure 4.**
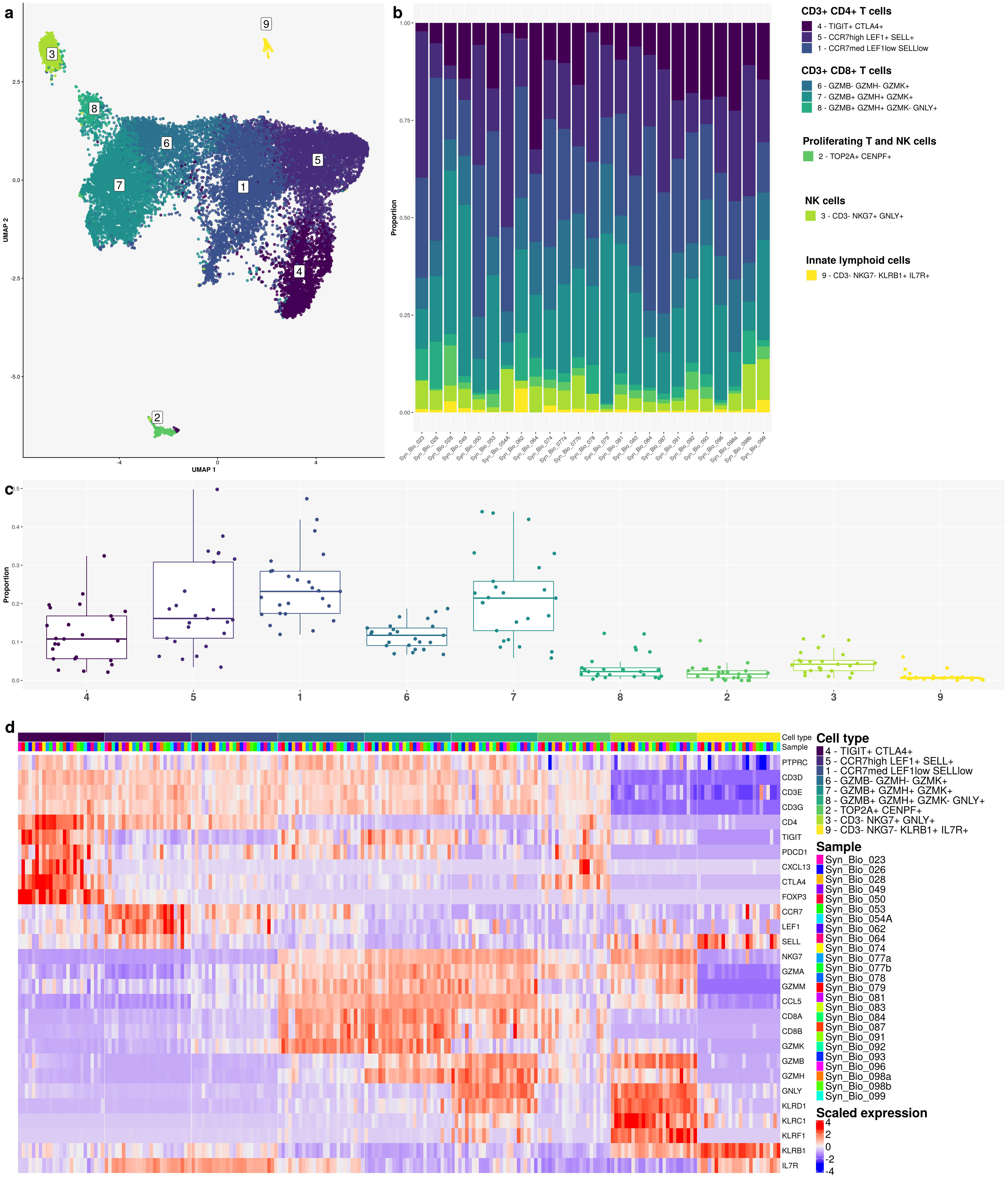
Subcluster analysis of synovial T cell, natural killer (NK) cell and innate lymphoid cell (ILC) populations in the integrated synovial single cell map datasets from 25 fresh-dissociated synovial biopsies of patients with inflammatory arthritis. **a)** UMAP with annotated T cell, NK cell and ILC clusters coloured by cell cluster, **b)** a bar plot of T, NK and ILC cell cluster abundances across synovial biopsies of patients with inflammatory arthritis coloured by cell cluster (right). **c)** the proportions of the nine cell clusters **d)** the heatmap of the top T cell, NK cell and ILC marker genes.

### Synovial fibroblast heterogeneity in freshly dissociated human synovium

Through our analysis, we identified seven COL1A1+ synovial fibroblast clusters (**Fig. 5a**) and further classified these cells based on their expression of the lining (PRG4) and sublining (THY1) fibroblast markers ^16, 21, 23^ (**Fig. 5a, Suppl. Fig. 9a**). Overall, the abundance of synovial fibroblast clusters varied significantly across patient synovia (**Fig. 5b, c**). The seven synovial fibroblast clusters represented lining PRG4 ^high^ THY1 ^low^ (cluster 3) synovial fibroblasts, transitional PRG4 ^med^ THY1 ^med/high^ synovial fibroblasts (clusters 4, 5, 7) and PRG4 ^low^ THY1 ^high^ sublining synovial fibroblasts (clusters 1, 2, 6) (**Fig. 5a, Suppl. Fig. 9a**). The proliferating TOP2A+ CENPF+ synovial fibroblasts co-clustered with cluster 4 fibroblasts and contained both PRG4^high^ and THY+ cells (**Suppl. Fig. 9b**). The PRG4 ^high^ lining synovial fibroblasts were previously reported as enriched in osteoarthritis synovia ^16^ and associated with matrix-degrading activities^20, 21^, while HLA-DRA+ sublining cells were found abundant in leukocyte-rich RA^16^.

**Figure 5.**
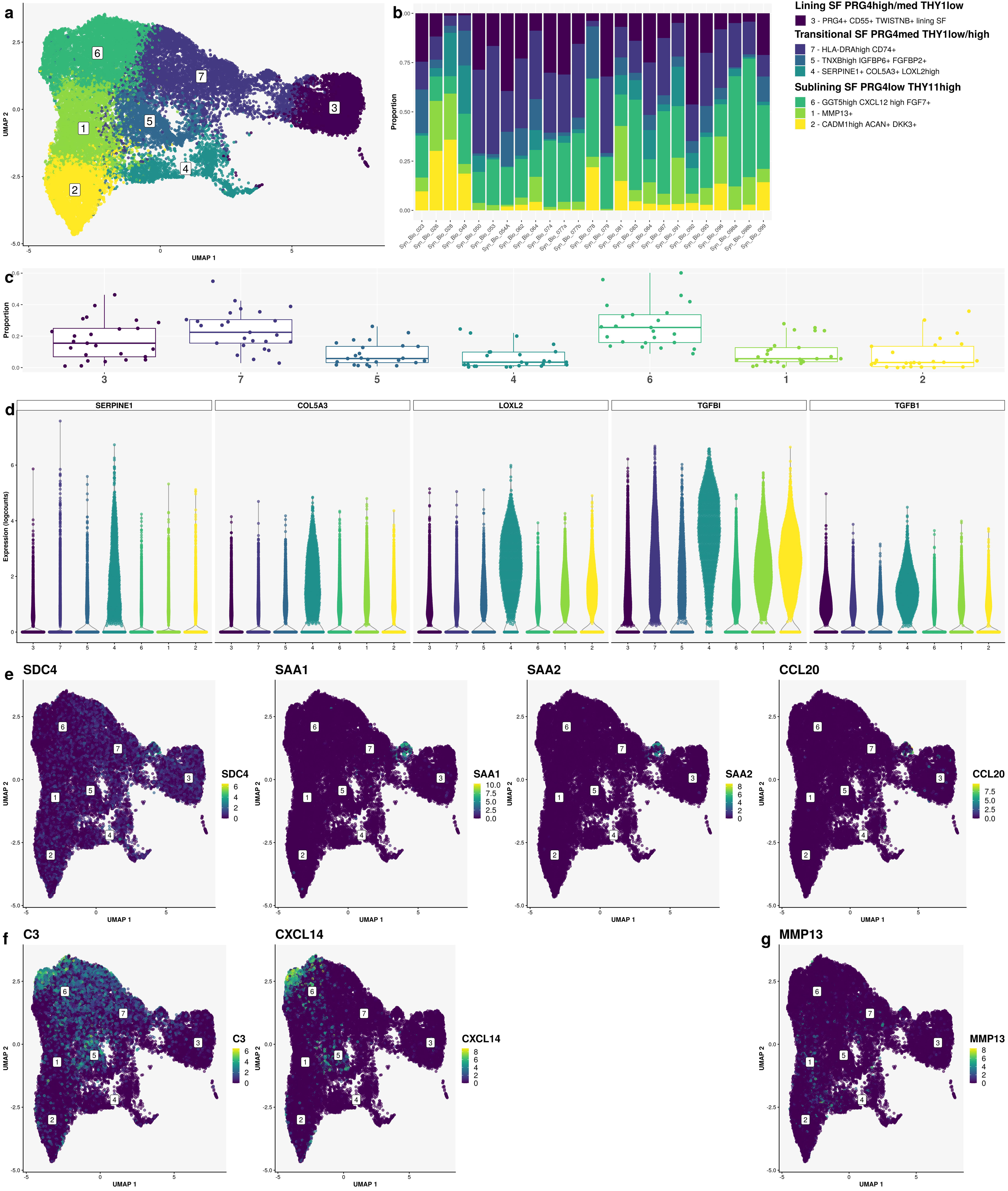
Subcluster analysis of synovial fibroblast population within the integrated synovial single-cell map dataset of inflammatory arthritis from 25 fresh-dissociated synovial biopsies of patients with inflammatory arthritis. **a)** UMAP with annotated synovial fibroblast clusters coloured by cell cluster. **b)** A bar plot of synovial fibroblast cluster abundances across synovial biopsy samples of patients with inflammatory arthritis coloured by cell cluster. **c)** The proportions of the seven synovial fibroblast clusters across 25 synovial samples from patients with inflammatory arthritis. **d)** Violin plots showing the top enriched matrix-remodelling genes (*LOXL2, TGFBI, TGFB1*), expressed by the SERPINE1+ COL5A3+ synovial fibroblasts (cluster 4). **e)** Violin plots showing the top enriched genes, expressed by the SOD2 ^high^ synovial fibroblast subpopulation within cluster 7, including *SDC4* and proinflammatory genes (*SAA1, SAA2, CCL20*). **f)** UMAPs demonstrating C3 and CXCL14 mRNA expression in synovial fibroblasts, identifying a small population of C3^high^ CXCL14^high^ cells within the sublining GGT5^high^ CXCL12^high^ synovial fibroblast subset (cluster 6). **g)** UMAP demonstrating MMP13 mRNA expression in synovial fibroblasts in cluster 1.

To further understand the synovial fibroblast transcriptional diversity, we analysed the expression of known fibroblast subset marker genes alongside top enriched genes between the seven fibroblast clusters (**Suppl. Fig. 10**). The lining fibroblasts (cluster 3) expressed CD55, ITGB8, MMP3, MMP1 and TWISTNB^53^ transcripts (**Suppl. Fig. 10**). These cells were GAS6 ^neg/low^ (**Suppl. Fig. 10**) as previously reported ^16, 18^. A subset of lining fibroblasts was enriched for CLIC5+ and HBEGF+ expression (**Suppl. Fig. 11a**). The cluster 4 represented SERPINE1+ COL5A3high fibroblasts, characterized by an extracellular matrix remodelling gene signature (*LOXL2, TGFBI, TGFB1*) (**Fig. 5d, Suppl. Fig. 10**). The HLA-DRA ^high^ cells (cluster 7) expressed transcripts linked to MHCII class antigen presentation (HLA-DRA, HLA-DRB1, HLA-DPB1, CD74, **Suppl. Fig. 11b**)^16^. HLA-DR and CD74 transcripts were expressed also in lining synovial fibroblasts. The cluster 7 contained a small population of HLA-DRA ^neg/low^ SOD2^high^ fibroblasts expressing syndecan 4 (*SDC4*) and proinflammatory genes *SAA1* and *SAA2* genes (**Fig. 5e**). SDC4 was previously reported to be prominently expressed in synovial lining cells in RA synovia^54^. Cluster 5 included TNXB ^high^ IGFBP6+ FGFBP2+ cells, expressing DKK3, CD34 and DPP4 transcripts (**Suppl. Fig. 10**). Among the sublining fibroblast clusters, GGT5 ^high^ CXCL12^high^ FGF7+ fibroblasts (cluster 6) contained NOTCH3-expressing cells, IL6-expressing cells (**Suppl. Figs. 10, 11c**) and a small population of C3 ^high^ CXCL14 ^high^ fibroblasts (**Fig. 5f, Suppl. Fig. 10**). NOTCH3 and GGT5 expression were previously associated with periarterial localization of synovial fibroblast^23^. Synovial fibroblasts in cluster 1 were specifically enriched in MMP13 (**Fig. 5a, g, Suppl. Fig. 10**); their transcriptome, however overlapped significantly with cluster 6 cells. Finally, cluster 2 cells represented CADM1^high^ ACAN+ DKK3+ sublining fibroblasts (**Fig. 5a, Suppl. Fig. 10**). Deconvolution analysis of synovial RNA-seq data demonstrated enrichment of DKK3+ synovial fibroblasts in patients with drug-refractory RA^55^.

### Synovial scRNA-seq analysis broadens gene signatures of synovial macrophage subsets

The synovial macrophage and DC cluster contained 35’659 CD45+ single cells distributed into CD14^low/neg^ CD64 ^low/neg^ CD11b^low/neg^ DC and CD14+ CD64+ CD11b+ macrophage clusters (**Fig. 6a, Suppl. Fig. 12**). The heatmap in **Suppl. Fig. 12** shows key marker and cluster-enriched genes expressed in macrophage and DC clusters.

**Figure 6:**
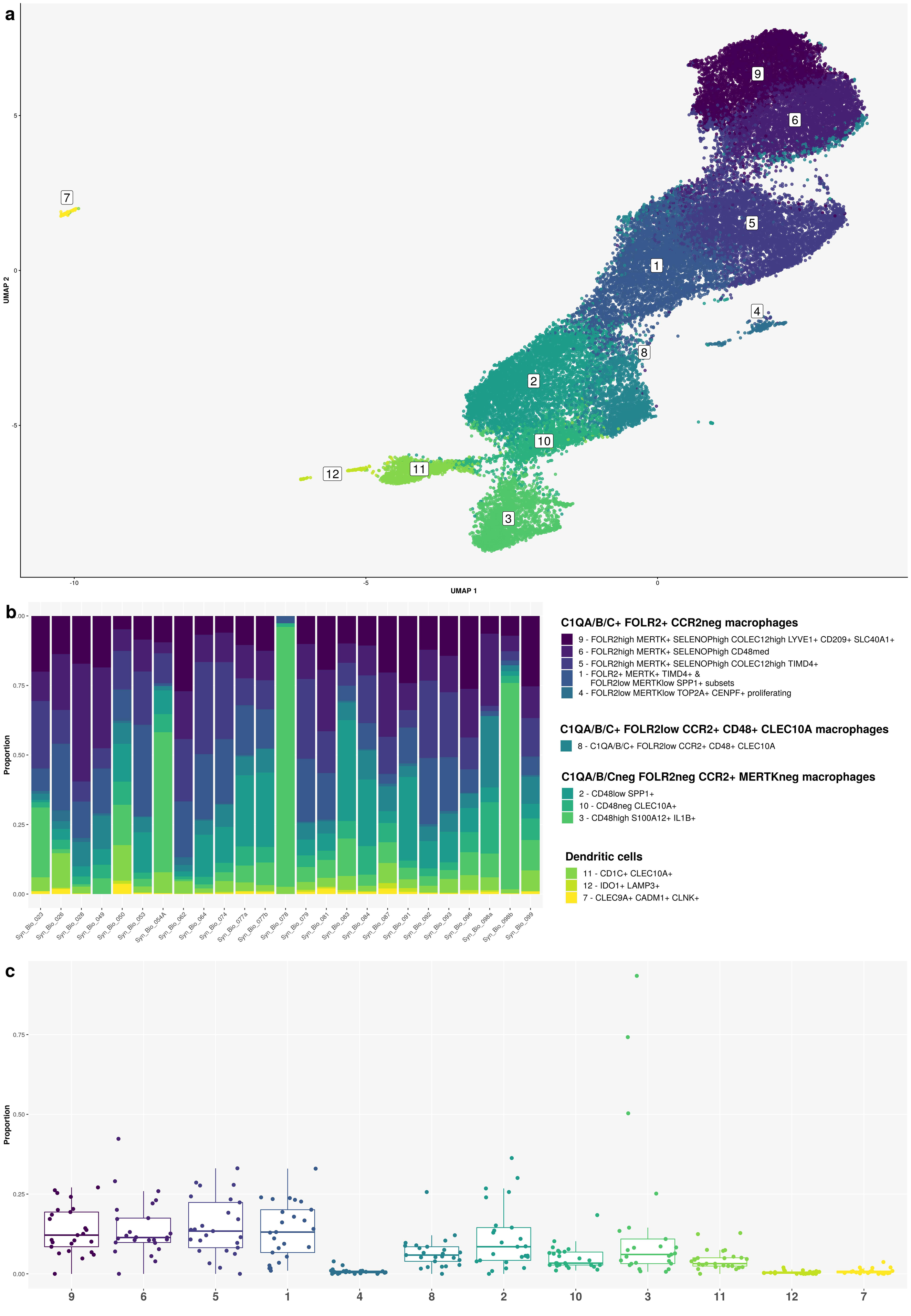
Subcluster analysis of synovial macrophage and dendritic cell (DC) populations in the integrated synovial single-cell map dataset from 25 fresh-dissociated synovial biopsies of patients with inflammatory arthritis. **a)** UMAP of 3 dendritic cell (DC) and nine synovial macrophage clusters coloured by cell cluster. **b)** Bar plots of relative abundances of three DC and nine synovial macrophage clusters across 25 patient samples coloured by cell cluster. **c)** The proportion of three DC and nine macrophage clusters across 25 synovial tissue samples from patients with inflammatory arthritis.

DC clusters included CD1C+ FCER1A+ CLEC10A+ DCs (cluster 11), CD83+ LAMP3+ IDO1+ DCs (cluster 12) and CLEC9A+ CADM1+ CLNK+ DCs (cluster 7) (**Fig. 6a, Suppl. Fig. 12**). We annotated the nine CD14+ CD64+ CD11b+ macrophage clusters based on their expression of CCR2, FOLR2 and complement chain 1q (C1QA, C1QB, C1QC) genes (**Figure 6a, Suppl. Fig 12**). The expression of CCR2 and FOLR2 was previously linked to tissue-infiltrating and tissue-resident macrophage subsets, respectively ^56^. Synovial macrophages grouped into C1QA/B/C+ FOLR2+ CCR2^neg^ (clusters 1, 4-6, 9), C1QA/B/C^neg^ FOLR2^neg^ CCR2+ (clusters 2, 3, 10) and C1QA-C+ FOLR2^low^ CCR2+ (cluster 8) clusters (**Suppl. Fig 12**). Macrophages in cluster 8 were TREM2+ MRC1+ MERTK^low^ and expressed CD48 and CLEC10A genes (**Suppl. Fig 12**).

The C1QA/B/C^neg^ CCR2+ macrophages comprised TREM2+ CD48^low^ SPP1+ (cluster 2), TREM2^neg^ CD48^high^ S100A12+ PLAC8+ SELL+ CD52 ^high^ (cluster 3) and TREM2 ^neg^ CD48+ CLEC10A+ (cluster 10) subsets (**Fig. 6a, Suppl. Fig. 12**). These cells were MERTK ^neg^ MRC1 ^low/neg^ and CD163 ^low^. SPP1+ macrophages were reported as enriched in RA synovitis ^18^ and bronchoalveolar lavage from patients with severe COVID ^52^. Furthermore, SPP1-expressing hepatic lipid-associated macrophages were shown to derive from bone marrow, replacing Kupffer cells (KCs) in metabolic-associated fatty liver disease ^57^. Genes linked to proinflammatory (*CD114, IL1B, S100A8, S100A9, FCAR*) and host defence (*FCN1, PPIF, CD93*) functions were highly expressed in cluster 3 cells (**Suppl. Fig. 12**).

The C1QA/B/C+ FOLR2+ CCR2-synovial macrophages segregated into FOLR2 ^high^ MERTK+ CD163^high^ SELENOP ^high^ (clusters 5, 6, 9), proliferating FOLR2 ^low^ MERTK^low^ TOP2A+ CENPF+ (cluster 4) and cluster 1 subsets, containing FOLR2 ^low^ MERTK^low^ SPP1+ and FOLR2+ MERTK+ TIMD4+ macrophage populations (**Figure 6a, Suppl Fig 12**). Macrophages in clusters 1, 4-6 and 9 were TREM2+; TREM2 has been linked to regenerative wound healing^58^ and pro-tumorigenic macrophage functions ^59^. They expressed the genes encoding *CMKLR1*, the receptor for resolvin E^60^, and *SLCO2B1* transporter, associated with macrophage identity ^61^. In addition to being enriched for complement component-encoding transcripts (*C1QA, C1QB, C1QC*), these cells expressed genes encoding scavenger receptors (*CD206, STAB1* ^62^, *CD163*), molecules involved in efferocytosis (*MERTK, LGMN*^63^) and cholesterol/lipid trafficking (APOE) (**Suppl. Fig. 12**), pointing to their tissue clearing functions. Looking closer at FOLR2 ^high^ MERTK+ CD163 ^high^ SELENOP^high^ macrophages (clusters 5, 6, 9), we identified COLEC12 ^high^ TIMD4+ macrophages (cluster 5), CD48^med^ macrophages (cluster 6) and COLEC12 ^high^ LYVE1+ macrophages (cluster 9) (**Fig. 6a, Suppl. Fig. 12**). TIMD4 and LYVE1 were associated with tissue residency ^64^ and the perivascular location ^65^, respectively. A scavenger receptor collectin family member 12 (COLEC12)^66^ was shown to be involved in the clearance of microbes, oxLDL ^67^ and myelin^68^ and acted as a high-affinity ligand for the collagen domain binding receptor LAIR1 ^69^. Studies in mice showed that LAIR1 controls homeostatic and anti-tumour functions of monocytes and interstitial macrophages in the lungs via stromal sensing ^69^. COLEC12+ tissue-resident macrophages in clusters 5 and 9 were specifically enriched for CCL18, CCL13 and NUPR1 (**Suppl. Fig. 12**), the regulators of lymphoid/myeloid cell trafficking ^70, 71^ and ferroptosis ^72^, respectively. Finally, LYVE1+ macrophages (cluster 9) expressed CD209, while being enriched in transcripts associated with coagulation (F13A1 ^73^), tissue adaptation to stress (IGF1 ^74^), suppression of inflammatory genes (EGR1^75^) and iron recycling (SLC40A1^76, 77^) (**Suppl. Fig. 12**). Like other synovial cell types, the abundances of DC and macrophage subsets varied significantly across patient synovia (**Fig. 6b, c**).

### Endothelial cell diversity in freshly dissociated human synovium in inflammatory arthritis

Our synovial scRNA-seq dataset contained 9395 PECAM+ synovial EC profiles forming eight EC clusters (**Fig. 7a**). We identified pan-endothelial and subtype-specific EC markers ^78^ and genes for the core EC functions (**Suppl. Fig. 13**). The eight EC clusters included LAPTM5+ PROX1+ LYVE1+ CCL21+ lymphatic ECs (cluster 4), proliferating TOP2A+ CENPF+ MKI67+ ECs (cluster 7), ACKR1+ venous ECs (clusters 1, 3, 5, 8), SPP1+ KDR+ capillary ECs (cluster 6) and GJA4+ CLDN5+ arterial ECs (cluster 2) (**Fig. 7a, Suppl Fig. 13**). The EC clustering UMAP demonstrates the principal separation of synovial ECs into vascular and lymphatic clusters (dimension 1), while dimensions 2 shows the core structure of tissue vasculature, starting with arterial, transiting into capillary and ending in venous vessel networks (**Fig. 7A**). Venous ACKR1+ ECs represented the largest synovial EC population (**Fig. 7a**). Among them, we identified ACKR1^med^ VWF^med^ KDR^low^ SPARC^high^ CLU^neg^ ECs (cluster 3) sharing capillary and venous gene expression signatures (**Suppl. Fig. 13**), suggesting a transitional phenotype. The ACKR1 ^high^ VWF^high^ KDR^neg^ CLU+ ECs (clusters 1, 5, 8) expressed IL1R1 transcripts (IL1B response) and were enriched for HLA-DR and HLA-DP transcripts (antigen presentation). Clusters 5 and 8 were additionally IL6+ SOD2+ SOCS3 ^high^ and expressed IRF1 (interferon response). ICAM1 (leukocyte trans-endothelial migration) and SELE (neutrophil adhesion) were enriched in clusters 5 and 3 cells, while *CCL2* and *TNFAIP3* genes were highest expressed in cluster 5 venous ECs (**Suppl. Fig. 13**). SPP1+ KDR+ capillary ECs (cluster 6) were strongly enriched for transcripts encoding basement membrane-associated collagens (COL4A1, COL4A2, COL15A1) ^79^ and plasmalemma vesicle associated protein (PLVAP) (**Suppl. Fig. 13**), a regulator of microvascular permeability and diaphragm formation within endothelial fenestrae ^80^. Finally, arterial ECs (cluster 2) were characterized by the expression of tight (*CLDN5*) and gap (*GJA4*) junction^81, 82^ genes as well as genes involved in endothelial barrier regulation (*RHOB*)^83^, lymphatic vascular patterning (*SEMA3G*)^84^, regulation of TGFB activity (*LTBP4*), lymphocyte trafficking (*ICAM2*) and chemotaxis (*CXCL12*) (**Suppl. Fig. 13**). The eight EC clusters were detectable in all patient samples, their abundancies, however, varied significantly across patient synovia (**Fig. 7b, c**).

**Figure 7:**
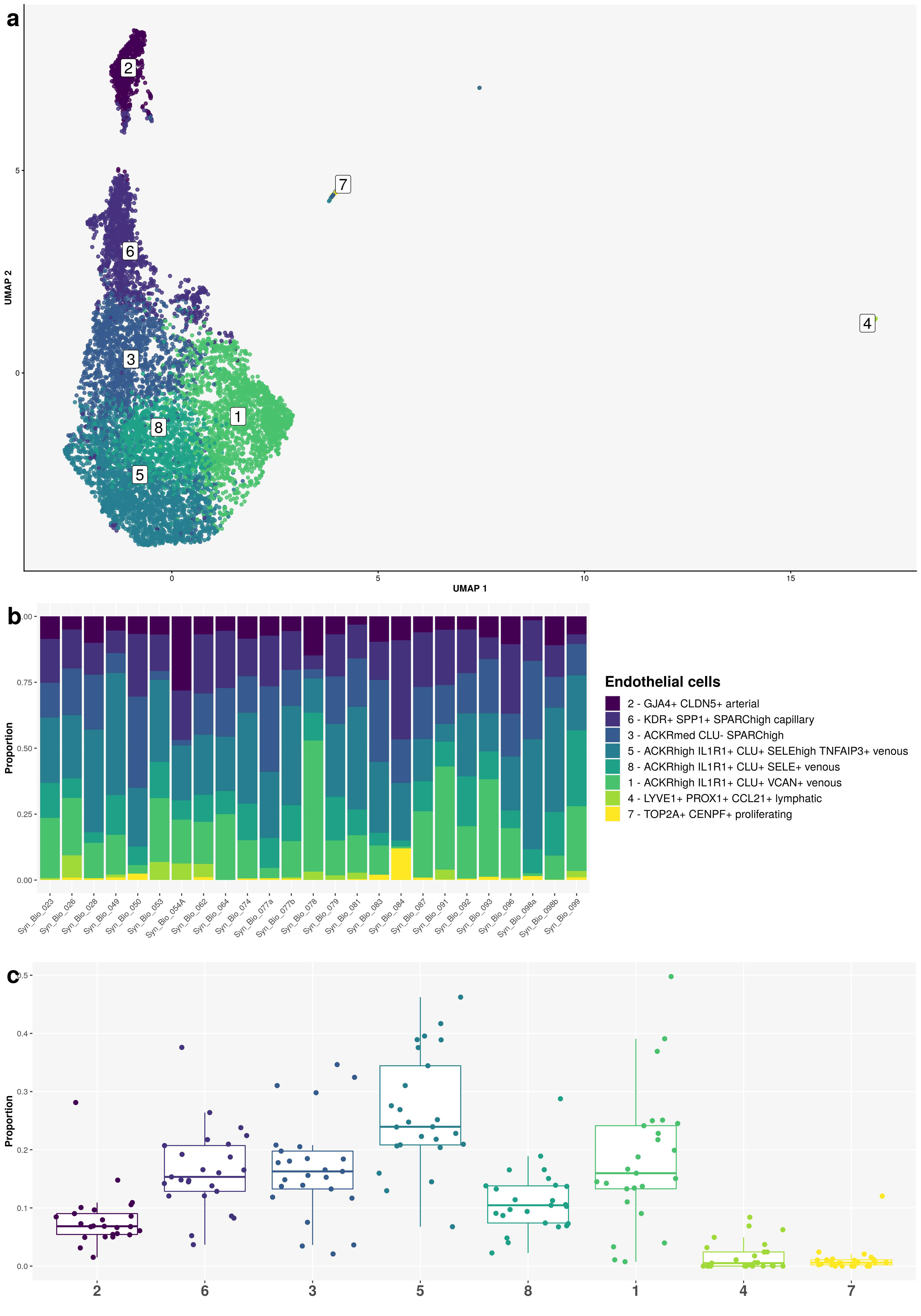
Subcluster analysis of synovial endothelial cell (EC) population in the integrated synovial single-cell map dataset from 25 fresh-dissociated synovial biopsies of patients with inflammatory arthritis. **a)** UMAP of eight synovial EC clusters coloured by EC cluster. **b)** Barplots of relative abundances of eight EC clusters across 25 synovial samples coloured by EC cluster. **c)** The proportion of the eight EC clusters across 25 synovial tissue samples from patients with inflammatory arthritis.

### A reference single-cell map of human synovium in inflammatory arthritis

In the next step, we merged the cell annotations from the main cell types (**Suppl. Fig. 6a**) and their subclusters (**Figs. 4-7**), identified in the integrated scRNA-seq dataset from 25 freshly dissociated synovia of patients with inflammatory arthritis. These annotations represented a detailed scRNA-seq reference map of the fresh-dissociated human synovial cells in inflammatory arthritis (**Fig. 8**).

**Figure 8.**
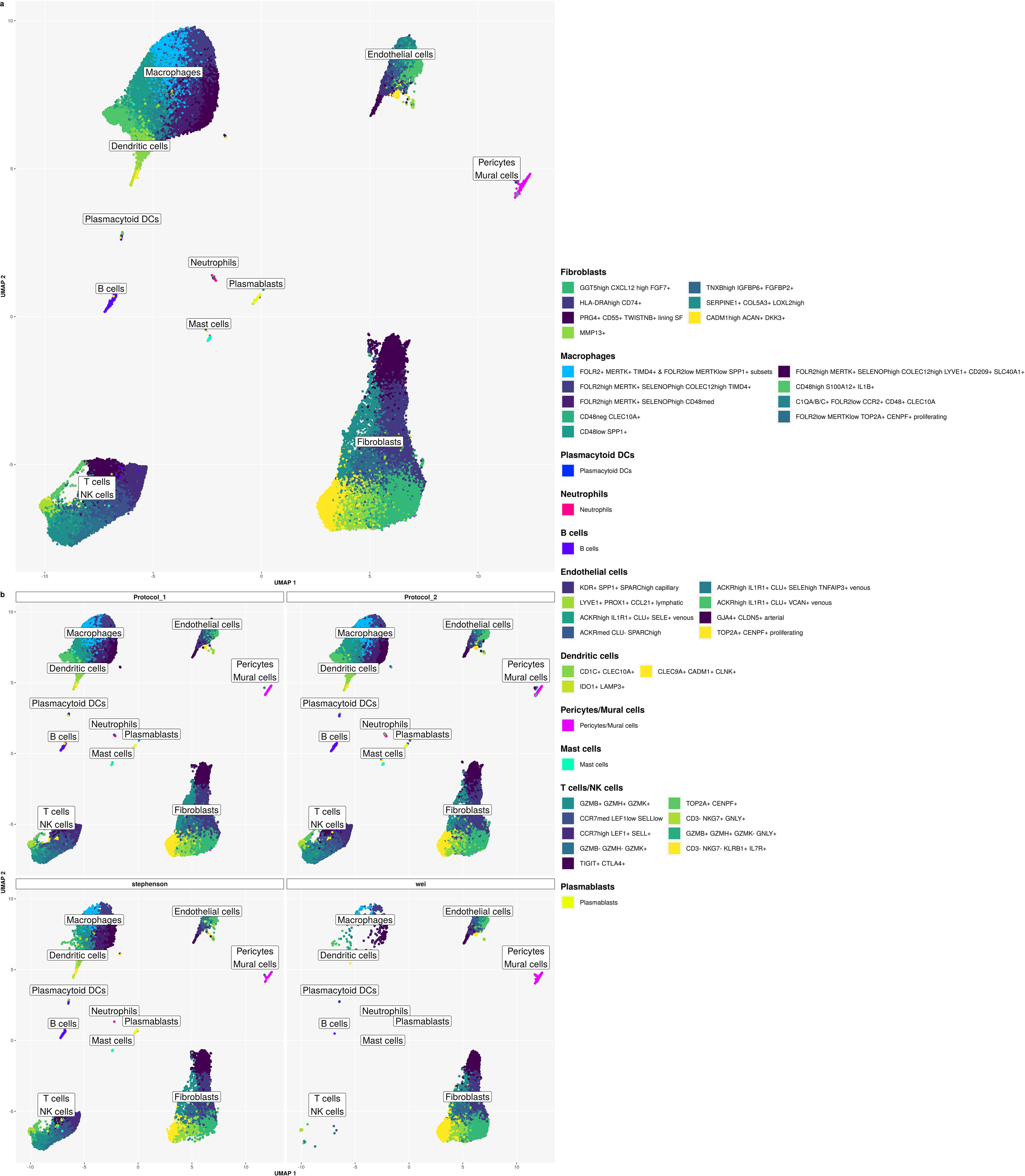
Single-cell reference map of the fresh-dissociated human synovium in inflammatory arthritis. A UMAP of annotated 12 lymphoid, 14 myeloid and 16 stromal cell synovial cell clusters and subclusters showing extensive cellular heterogeneity of the human synovium in inflammatory arthritis. Cluster annotations are based on the analysis of the main cell types and the subcluster analysis of synovial T-cells/natural killer (NK) cells/innate lymphoid cells (ILC), fibroblast, macrophage, dendritic cell (DC) and endothelial cell populations. The integrated scRNA-seq data are derived from 25 synovial tissue samples from patients with different types of inflammatory arthritis (see Methods for details).

### Our synovial scRNA-seq cell map serves as a reference annotation resource to map synovial cell types in published synovial scRNA-seq datasets

As the final step in our analysis, we integrated our synovial scRNA-seq dataset (**Fig. 8**) with publicly available synovial scRNA-seq data. We utilized our synovial cell map as a reference (**Fig. 8**) to annotate cell types across the integrated datasets. We selected a dataset from Stephenson at colleages^17^ generated from unsorted fresh dissociated synovial cells, alongside a dataset from Wei K and colleagues^23^ containing a large set of scRNA-seq synovial endothelial cell profiles, excluded from most other synovial scRNA-seq datasets. Using our synovial reference cell map, we successfully annotated the major synovial cell types and their subsets in the integrated (our, Stephenson and Wei) dataset (**Fig. 9a, b, Suppl Fig. 14a**). Stephenson’s dataset included all synovial cell types, including a minor synovial neutrophil population (**Fig. 9b, Suppl. Fig. 14b, c**). Wei’s dataset comprised primarily synovial fibroblast, mural cells/pericytes and endothelial cell subsets (**Fig. 9b, Suppl. Fig. 14b, c**), which were selectively sorted out from synovial tissue prior to scRNA -seq analysis. Abundance of main cell clusters varied significantly across patients in all datasets (**Suppl. Fig. 14b, c**).

**Figure 9.** Utilization of our synovial scRNA-seq reference dataset for annotating publicly available synovial scRNA-seq datasets. Lower dimensional representation (UMAP) of our scRNA-seq dataset (“Protocol 1” and “Protocol 2”) integrated with two publicly available datasets (Stephenson W and colleagues ^17^ and Wei K et colleagues ^23^). The colour represents the cell type as determined by either manual annotation (“Protocol 1” and “Protocol 2”) or label transfer (“Stephenson” and “Wei”) (see Methods). Annotated are 12 lymphoid, 14 myeloid and 16 stromal cell synovial cell clusters and subclusters (see Fig. 8 for details). **a)** Integrated data from our, Wei K et al. ^23^ and Stephenson W et al. ^17^ studies combined in one graph, **b)** UMAPs with scRNA-seq data split by protocol and study.

## Discussion

Integrative single-cell omics is quickly expanding our understanding of unique and shared cell types across human tissues in health and disease ^85^. These analyses require isolation of highly viable cells either from fresh or cryopreserved human tissues, the latter often necessitating viable cell sorting.

Fresh tissue dissociation facilitates less biased cell interrogation with valuable insights into the biology and heterogeneity of cryopreservation-sensitive cell types like neutrophils. Analyzing unsorted synovial cells from prospectively collected fresh tissues relies strongly on the stable isolation of highly viable cells to prevent sample loss. Here, we utilized the protocol of Donlin et al., optimized for the dissociation of cryopreserved synovia ^22^, and refined it for the dissociation of small fresh human synovial biopsies. We enhanced the release of synovial cells by gentle tissue massaging and minimized cell loss by optimizing the washing steps and volumes of reaction mixes. These protocol refinements were associated with consistent isolation of good yield of viable synovial cells, thereby overcoming significant sample loss, associated with the original protocol. Overall, we established a reliable dissociation method for fresh, prospectively collected synovial biopsies that facilitates scRNA-seq studies on unsorted synovial cells. This method complements the original protocol by Donlin and colleagues, which is optimized for multicentric omics studies on cryopreserved synovia^22^.

We detected a minor population of IFITM2 ^high^ neutrophils in scRNA-seq data from freshly dissociated synovia. A paucity of detected neutrophils was in line with histological knowledge about inflamed RA synovium, where neutrophils are commonly scarcely present. We detected neutrophils mainly in samples from protocol 2 donors, as evidenced by histology and scRNA-seq analyses. The overlap of both methods in neutrophil detection was 50%, which could be attributed to neutrophil scarcity and their focal distribution within and across biopsy fragments. The integrative scRNA-seq analysis of 25 synovial tissue samples further inferred that synovial neutrophil detection was primarily donor-dependent and not protocol-specific (**Suppl. Fig. 6b**). In general, sequencing of a high number of total non-sorted synovial cells and dissociating synovium fresh could facilitate neutrophil detection in our study irrespective of the protocol used. To further explore neutrophil abundances in human synovia, we aimed to integrate in-house and publicly available human synovial datasets. The short lifespan and intrinsically small library sizes of human neutrophils ^86^ have challenged the analysis of neutrophil abundances in publicly available synovial scRNA-seq data. Additionally, most published synovial scRNA-seq datasets were created on pre-sorted synovial cells, excluding neutrophils^16, 18, 23^. We integrated our dataset with two publicly available datasets, including data from 20’000 fresh dissociated unsorted synovial cells from Stephenson and colleagues ^17^ available in the pre-processed form of the filtered count matrices. This integrative analysis demonstrated the presence of minor neutrophil populations in our and Stephenson’s data. No neutrophils were detected in Wei’s data ^23^, created on sorted CD45-CD235-synovial stromal cells from cryopreserved synovial tissues.

Our study reproduced many previously described synovial fibroblast, myeloid and lymphoid transcriptional cell states ^16–18, 23^, while expanding synovial cell heterogeneity to neutrophils. Furthermore, we identified a small population of SDC4 + synovial fibroblasts and SERPINE1+ COL5A3+ synovial fibroblasts, while broadening the transcriptional characterization of tissue-resident COLEC12 ^high^ synovial macrophages. SDC4 has been closely linked to synovial fibroblast attachment/invasion into articular cartilage and cartilage breakdown ^54, 87, 88^, while COLEC12 and SLC40A1 were associated with macrophage extracellular matrix sensing ^69^ and regulation of iron homeostasis ^76, 77^. We observed a minor subset of C3 ^high^ CXCL14 ^high^ cells within the sublining GGT5 ^high^ CXCL12 ^high^ synovial fibroblast cluster. Notably, synovial fibroblasts were recently shown to drive the local inflammatory tissue priming in preclinical models of arthritis in a C3-dependent manner; the primed fibroblast appeared to upregulate both C3 and CXCL14 mRNA expression ^89^. Future phenotypic and functional experiments should confirm whether gene signatures and annotated cell clusters also translate into specific synovial cell functions.

Differential enrichment of distinct cell types across patient synovia might drive different pathogenic pathways and contribute to the characteristic interpatient variability in therapeutic responses to currently prescribed disease-modifying antirheumatic drugs ^55^. Our in-depth analysis of synovial EC diversity in inflammatory arthritis demonstrated the shared contribution of GGT5+ synovial fibroblasts and ACKR1^high^ SELE ^high^ TNFAIP3+ venous EC subsets to the synovial production of IL-6. The variable abundances of IL6-producing cell subsets across patient synovia may shape the differential therapeutic responses of arthritis patients to anti-IL6 therapies or JAK inhibitors. Future studies involving larger stratified patient cohorts will be essential to link synovial cell composition to variation in therapeutic response. Furthermore, such studies might contribute to the cell subtype-based and molecular pathway-based patient stratification across and within different clinical types of inflammatory arthritis^90^.

In conclusion, we refined the synovial dissociation protocol for prospectively collected fresh synovial biopsies and generated an extensive reference single-cell resource of freshly dissociated human synovium in inflammatory arthritis. Our synovial cell annotations were efficiently used to annotate published synovial scRNA-seq datasets, thereby serving as a reference human synovium map. This study not only diversifies the knowledge about synovial cell composition in inflammatory arthritis but could also facilitate future prospective single-cell omics studies on fresh human synovium.

## Acknowledgements

We thank all the patients who donated synovial biopsies for the realization of this study and Benvinda Campos Henriques for excellent technical support.

## Author contributions

S.G.E, R.G., M.D.R and M.F.B conceived the study, S.G.E, M.H., T.K, N.I., B.B., M.F.B, performed experiments, R.G., M.D.R conducted a bioinformatic analysis of scRNA-seq data, R.M., K.B. recruited the patients and collected clinical data, R.M. performed the ultrasound-guided synovial biopsy, Ž.R., M.T., S.Č., O.D., C.O., S.S.S, M.F.B were involved in study organization, C.P. performed immunohistology, C.P. and R.M. interpreted immunohistology data, S.G.E, R.G., M.D.R. and M.F.B analyzed and interpreted scRNA-seq data, S.G.E, R.G., M.H., M.D.R. and M.F.B wrote the paper, all authors contributed to critical discussion and final drafting of the article, and agree with the paper content.

## Supplementary Figures

**Supplementary Figure 1**. Quality control of integrative protocol scRNA-seq analysis of 18 human synovial biopsies of patients with inflammatory arthritis. **a)** The number of genes vs. total counts per sample coloured by filter (left) and percentage mitochondrial counts (right). **b)** The number of cells per sample (left) and per protocol (right), coloured by the number of genes (top) and by the number of genes in at least 1% of cells (bottom). **c)** Distribution of counts per protocol (left) and per sample (right). **d)** Distribution of the number of genes per protocol (left) and per sample (right).

**Supplementary Figure 2.** UMAPs of integrated scRNA-seq data from integrative protocol analysis (18 synovial tissue biopsies of patients with inflammatory arthritis), colour by top marker genes of major synovial cell populations including **a)** T cells, B cells and plasmablasts; **b)** macrophages and plasmacytoid dendritic cells; **c)** myeloid dendritic cells, mast cells and neutrophils; **d)** endothelial cells, pericytes/mural cells and synovial fibroblasts.

**Supplementary Figure 3.** UMAPs of integrated scRNA-seq data from integrative protocol analysis. Data from 18 synovial tissue biopsies of patients with inflammatory arthritis. **a)** T cells/NK cells, colour by key marker genes *CD3*, *CD4*, *CD8*, *NKG7*. **b)** Macrophages color by key marker genes *MERTK, TREM2, CD206/MRC1, IL1B, CD4* 8. **c)** Endothelial cells, colour by key marker genes *LYVE1, CCL21, VWF*. **d**) Synovial fibroblasts, colour by key marker genes *PRG4, THY1, GGT5, CXCL12, DKK3*.

**Supplementary Figure 4:** The heatmap of major neutrophil genes, detected in the integrative protocol scRNA-seq analysis, data from 18 synovial tissue biopsies of patients with inflammatory arthritis. Expressions are aggregated by sample and cell type.

**Supplementary Figure 5.** Quality control of scRNA-seq analysis of the single cell reference map of fresh human synovium from 25 synovial biopsies of patients with inflammatory arthritis. **a)** A cell level summary of the total number of counts, the number of detected genes and the percentage of mitochondrial counts. The number of genes vs. total counts per sample coloured by filter (left) and percentage mitochondrial counts (right). **b)** Sample summary statistics with the number of cells and number of detected genes after filtering of low-quality cells. The number of cells per sample is coloured by the number of genes (top) and by the number of genes in at least 1% of cells (bottom). **c)** Distribution of counts per sample. **d)** Distribution of the number of genes per synovial tissue sample.

**Supplementary Figure 6.** Integrated scRNA-seq dataset from 25 synovial tissue samples from patients with inflammatory arthritis (see Methods for details). **a)** UMAP of annotated main synovial cell populations coloured by main cell type. **b)** Bar plots of relative abundances of main cell types per sample coloured by main cell type. **c)** The variability of the proportion of cell types across patient synovial tissues.

**Supplementary Figure 7.** A heatmap of top marker genes of the main synovial cell types identified in the integrated scRNA-seq dataset across 25 synovial tissue samples from patients with inflammatory arthritis (see Methods for details). Expressions are aggregated by sample and cell type.

**Supplementary Figure 8. a-d)** T cell and natural killer (NK) cell and innate lymphoid cell subclustering. The integrated scRNA-seq dataset consists of scrNA-seq profiles from 25 synovial tissue samples from patients with different types of inflammatory arthritis (see Methods for details). **a)** UMAPs showing the small population of proliferating TOP2A+ CENPF+ T cells (cluster 2). **b)** A heatmap of the cluster-enriched genes and marker genes in synovial T cell, NK cell and innate lymphoid clusters in patients with inflammatory arthritides. Expressions are aggregated by sample and cell type. **c)** UMAPs demonstrating the expression of *FOXP3, PCDC1* and *CXCL13* genes in the TIGIT+ CTLA4+ T cells (cluster 4). **d)** UMAPs demonstrating the expression of granzyme (*GZMK, GZMB, GZMH*) and granulysin (*GNLY*) genes in CD8+ T cells (clusters 6-8) and NK cells (cluster 3).

**Supplementary Figure 9. a-b)** Synovial fibroblast subclustering. The integrated dataset from 25 synovial biopsies of patients with inflammatory arthritis (see Methods for details) with **a)** violin plots showing the expression (log counts) of the lining marker PRG4 and the sublining marker THY1 across the seven synovial fibroblast clusters, and **b)** UMAPs showing the small population of proliferating TOP2A+ CENPF+ synovial fibroblasts, co-clustering with the cluster 4 SERPINE1+ COL5A3+ synovial fibroblasts (see also **Fig. 5a, d**).

**Supplementary Figure 10.** A heatmap of the top cluster genes and known markers of synovial fibroblast clusters detected in the synovium of patients with inflammatory arthritides. The integrated scRNA-seq dataset consists of 25 synovial tissue samples from patients with different types of inflammatory arthritis (see Methods for details). Expressions are aggregated by sample and cell type.

**Supplementary Figure 11. a-c)** Synovial fibroblast subclustering. The integrated dataset from 25 synovial biopsies of patients with inflammatory arthritis (see Methods for details) with **a)** UMAPs showing the higest enriched expression of the genes involved in MHCII class mediated antigen presentation in HLA-DRA^high^ synovial fibroblasts (cluster 7), and **b)** UMAPs showing the enriched expression of *IL6* and *NOTCH3* genes primarily in sublining GGT5+ synovial fibroblasts (clusters 6).

**Supplementary Figure 12.** Synovial macrophage and myeloid dendritic (DC) cell subclustering. A heatmap of the top cluster genes and known marker genes of synovial macrophage and DK subclusters detected in the integrated scRNA-seq dataset from 25 synovial tissue samples of patients with different types of inflammatory arthritis (see Methods for details). Expressions are aggregated by sample and cell type.

**Supplementary Figure 13.** Endothelial cell subclustering. Heatmap of the top cluster genes and known markers of synovial vascular and lymphatic endothelial cell subclusters detected in the integrated scRNA-seq dataset from 25 synovial tissue samples of patients with different types of inflammatory arthritis (see Methods for details). Expressions are aggregated by sample and cell type.

**Supplementary Figure 14. Integration of our synovial scRNA-seq from fresh synovium with publicly available synovial scRNA-seq datasets.** We integrated our scRNA-seq data from 25 synovial tissue samples of patients with inflammatory arthritis with published data from Stephenson W and colleagues^17^ and Wei K et colleagues ^23^ (see Methods for details). **a)** UMAP of annotated main synovial cell populations coloured by main cell type across the three studies. **b)** Bar plots of relative abundances of main cell types per sample coloured by main cell types across the 3 studies. **c)** The variability of the proportion of cell types across patient synovia in the 3 studies. Our data and data from Stephenson W et al. are derived from unsorted synovial cells, while Wei K et al dataset includes sorted CD45 ^neg^ CD235 ^neg^ synovial fibroblasts, pericytes/mural cells and synovial endothelial cells.

## Supplementary Tables

**Supplementary Table 1.** Detection of neutrophils in 18 synovial tissue biopsies included in integrative protocol analysis using histology and scRNA-seq analyses. For the detection of neutrophils in formalin-fixed, paraffin-embedded synovial biopsy fragments, tissue sections were labelled with CD15 antibodies (see Methods for details). In scRNA-seq data, the quantity of neutrophils was estimated by calculating the proportion of neutrophils per sample. Data pertain to 18 samples included in integrative protocol analysis.

| Pat.-ID. | Details on neutrophil histology, based on CD15 staining. | Proportion of neutrophils % | Histology. Neutrophil | ScRNA-seq. Neutrophil |
| --- | --- | --- | --- | --- |

|  |  | (scRNA-seq).* | detection. | detection. |
| --- | --- | --- | --- | --- |
| **SB-023 P1** | Focal, in addition fibrin associated neutrophils | 0.000173 | Y | y |
| **SB-026 P1** | Single fibrin associated neutrophils | 0 | Y | N |
| **SB-028 P1** | Fibrin associated neutrophils | 0 | Y | N |
| **SB-049 P1** | No CD15 staining | 0 | N | N |
| **SB-050 P1** | Focally interspersed neutrophils | 0 | Y | N |
| **SB-074 P1** | No CD15 staining | 0 | N | N |
| **SB-077A P2** | Up to 10 neutrophils per HPF (0.26mm<sup>2</sup>) | 0 | Y | N |
| **SB-077B P2** | 1 fragment with neutrophils, up to 20 per HPF (0.26mm<sup>2</sup>) | 0 | Y | N |
| **SB-079 P2** | <2 per HPF (0.26mm<sup>2</sup>) | 0 | Y | N |
| **SB-081 P2** | Up to 15 neutrophils per HPF (0.26mm<sup>2</sup>) | 0.004424 | Y | Y |
| **SB-083 P2** | Fibrin associated neutrophils | 0.007027 | Y | Y |
| **SB-084 P2** | Vessel associated neutrophils | 0.000757 | Y | Y |
| **SB-087 P2** | Focal florid inflammation with more than 35 neutrophils per HPF (0.26mm<sup>2</sup>) | 0.002684 | Y | Y |
| **SB-092 P2** | Focal small infiltration | 0 | Y | N |
| **SB-093 P2** | Predominantly fibrin associated neutrophils with transition into synovial membrane, up to 5 per HPF (0.26mm<sup>2</sup>) | 0.000206 | Y | Y |
| **SB-096 P2** | No CD15 staining | 0.001002 | N | Y |
| **SB-098a P2** | Up to 38 neutrophils per HPF (0.26mm<sup>2</sup>) | 0 | Y | N |
| **SB-098b P2** | Fibrin associated neutrophils | 0.056911 | Y | Y |

## Supplementary Methods

### Step-by-step synovial tissue dissociation protocol

#### A) Digest synovial tissue

In the first step of the protocol, fresh synovial tissue is washed to remove the potential sources of non-synovial cell contamination, minced into small fragments, and dissociated into synovial cell suspension using the combined enzymatic-mechanical tissue dissociation protocol. **CRITICAL**: Volumes of digestion mixes are refined to minimize cell loss.

##### Timing: 47 min

1. Wash synovial biopsy fragments in D-PBS.
  a. Transfer the biopsied tissue fragments with sterile forceps onto a wet membrane of a 70 μm cell strainer sitting in a well of a 6-well plate, filled with 4 ml PBS.
  b. Transfer the strainer with tissue to neighboring wells with fresh 4ml D-PBS. Repeat this washing step 2×. Keep the tissue wet. Washing removes potential cell contaminants that do not originate from tissue.
2. Mince synovial tissue.
  a. Transfer tissue fragments into a 250μl pre-warmed RPMI (glutamine, HEPES, no Abs, no FBS) medium droplet on a round bottom culture plate.
  b. Mince synovial tissue with a sterile scalpel into small tissue fragments (∼1mm). Keep the tissue wet. Prepare digestion mix.
  c. Transfer 500μl of the pre-warmed RPMI medium (glutamine, HEPES, no Abs, no FBS) into a conical 15ml polystyrene tube. Add 40μl of 2.5mg/ml Liberase TL, Roche to reach 100 μg/ml working Liberase TL concentration and 10μl of 10 mg/ml DNAse I, Roche to get 100 μg/ml working DNase I concentration.
  d. Transfer the minced tissue fragments with a wide-bore pipette tip, forceps or scalpel into the digestion mix and top the volume with RPMI (glutamine, HEPES, no Abs, no FBS) to a total of 1ml. Collect all tissue fragments, which might stick to the tip, scalpel, or forceps surfaces. Avoid using forceps with serrated surfaces to prevent excessive tissue sticking and tissue loss.
  e. Add magnetic stirrer(s) in the tube and close the tube safely.
3. Digest synovial tissue at 371Z°C in a mixed enzymatic-mechanical protocol.
  a. Transfer the tube with tissue fragments into a 371Z°C water bath on a magnetic holder, placed within the prewarmed oven (371Z°C). **NOTE**: The water bath facilitates keeping the suspension temperature at 371Z°C during tissue digestion.
  b. Digest tissue for 30 min at 371Z°C with a continuous magnetic stirring. **NOTE:** A combination of a ball-shaped and a cylinder-shaped stirrer facilitates tissue mixing during digestion.
  c. Fifteen minutes after starting the digestion, pass tissues gently through a 16G needle (10x) using a 1 ml syringe (mechanical dissociation). **CRITICAL**: The 16G needle may clog with tissue fragments; the built pressure may detach the needle, leading to tissue fragment/cell loss. The 1 ml syringe fits in the falcon tube, preventing the loss of cell suspension in case of needle detachment.
4. Stop enzymatic digestion.
  a. Add 200μl of FBS and 800μl of RPMI (glutamine, HEPES, no Abs, no FBS) to the digestion mix to reach the final 10% FCS concentration.

#### **B)** Enrich dissociated synovial cells

In this step, tissue debris and cell aggregates are removed to obtain a single-cell suspension of synovial cells.

**CRITICAL**: Synovial cell suspension is enriched by enhancing release of synovial cells from the digested tissue. Additional washes are introduced to minimize cell loss.

##### Timing: 16 min

1. Pre-wet the 40µm strainer with 2ml of RPMI medium (glutamine, HEPES, no Abs, 10% FBS) in the six-well plate. Filter the cell suspension through the 40µm strainer submerged into the medium using 1ml wide-bore pipette tips.
2. Wash the falcon tube with an additional 1ml of 10% RPMI to collect the remaining cells and filter through the pre-wet 40µm strainer into the well with filtered cell suspension.
3. Tissue debris and poorly disaggregated cell clumps will remain on the strainer. Using the syringe plunger head, gently press the synovial tissue fragments against the strainer to facilitate further cell release. **NOTE:** Collect any remaining drops of cell suspension from the bottom side of the strainer with a fresh pipette tip and add to the filtered cell suspension. **CRITICAL:** This step releases additional cells from the tissue.
4. Transfer the 5ml of the filtered cell suspension into a new 15ml Falcon tube.
5. Wash the well with an additional 5ml of 10% RPMI to collect the remaining cells.
6. Centrifuge the enriched filtered cell suspension at 300xg, RT, 10 min.

#### **C)** Lyse red blood cells

In this step, red blood cells are lysed, and synovial cells are washed.

##### Timing: 27 min

1. Lyse the red blood cells.
  a. During centrifugation, prepare 1× red cell lysis buffer (Roche) by mixing 0.5ml of 10x RBC Lysis Buffer in 4.5ml of nuclease-free H2O.
  b. Remove the supernatant and resuspend the pellet gently in 0.5ml RPMI containing 10% FCS.
  c. Add 5 ml of 1× RBC lysis buffer to the cells in 10% RPMI, vortex gently 5 sec, incubate 2 min RT.
  d. Centrifuge cell suspension at 300xg, RT, 10min. Remove the supernatant.
2. Wash the cells in D-PBS.
  a. Resuspend cells gently by flicking the pellet and then wash the cells with 10ml PBS. Centrifuge at 300xg RT, 10 min.
  b. Remove the supernatant.
  c. Centrifuge briefly at 300xg, RT to remove any residual solution.

#### **D)** Prepare single-cell suspension for scRNA-seq (10xGenomics)

In this step, the enriched cells are counted and checked for viability, potential remaining cell debris and aggregates are removed by additional filtering steps, and the suspension of single synovial cells is prepared for downstream applications (e.g., scRNA-seq).

##### Timing: 10 min

1. Prepare fresh 0.2% BSA D-PBS (4μl of 50mg/ml BSA per 100μl D-PBS).
2. Gently flick the pellet and resuspend the cells with wide-bore tips in 100-150μl in 0.2% BSA D-PBS. Keep the cells on ice.
3. Count the cells, determine cell viability and check for debris or cell clumps (e.g., with Luna-FL dual fluorescence cell counter, Logos BioSystems). If remaining debris or cell clumps, filter the cell suspension through the 35μm strainer into a 1.5ml Eppendorf tube. **CRITICAL:** This step helps to prevent the clogging of the 10xGenomics chips.
4. Adjust the cell number to 700 single synovial cells/μl. Dilute or concentrate the cell suspension in 0.2% BSA D-PBS accordingly. **NOTE:** 700 cells/μl in our experience minimizes the probability of chip clogging.
5. Keep the cells on ice until starting with the 10x Genomics protocol.

## Key Resource Table

| Reagents, commercial assays, consumables | SOURCE | IDENTIFIER |
| --- | --- | --- |
| Dulbecco's Phosphate Buffered Saline | Thermo Fisher Scientific | Cat#14190144 |
| RPMI 1640 medium, HEPES | Thermo Fisher Scientific | Cat#52400025 |
| Liberase TL | Roche | Cat#05401020001 |
| DNase I | Roche | Cat#11284932001 |
| Fetal Bovine Serum | Corning | Cat#35-079-CV |
| Red Blood Cell lysis solution | Miltenyi Biotec | Cat#130-094-183 |
| UltraPure™ DNase/RNase-free distilled water | Thermo Fisher Scientific | Cat#10977035 |
| UltraPure™ BSA (50 mg/ml) | Thermo Fisher Scientific | Cat#AM2616 |
| Acridine Orange PI | BioCat | F23002-LG |
| Chromium Controller & Next GEM accessory kit | 10x Genomics | Cat#1000204 |
| Chromium Next GEM Single Cell 3' GEM, library & gel bead kit v3.1, v3.0 | 10x Genomics | Cat#1000121 |
| Chromium Next GEM Chip G single cell kit | 10x Genomics | Cat#1000120 |
| Single Index Kit T Set A | 10x Genomics | Cat#1000213 |
| 6-well cell culture plate | TPP | Cat#92006 |
| Cell strainer (70 µm) | Corning | Cat#352350 |
| ART™ wide bore filtered pipette tips | Thermo Fisher Scientific | Cat#2069G |
| Tissue culture dish | TPP | Cat#93100 |
| Scalpel | B. Braun | Cat#5518067 |
| Tweezers |  |  |
| 15ml centrifuge tubes | TPP | Cat#91015 |
| Magnetic stirring bar – ball shaped, micro |  |  |
| Digital incubator 37°C |  |  |
| Magnetic stirrer/squid |  |  |
| Beaker (100ml) |  |  |
| 16-gauge blunt needle | Maybach | Cat#8001741 |
| 3 -piece syringes (1ml) | BD Medical | Cat#303172 |
| Cell strainer (40µm, 70µm) | Corning | Cat#352340 |
| Centrifuge |  |  |
| Luna-FL™ Dual Fluorescence counter | Logos Biosystems |  |

