## Supplementary Figures for "A reference single-cell map of freshly dissociated human synovium in inflammatory arthritis with an optimized dissociation protocol for prospective synovial biopsy collection"

### Slide 1
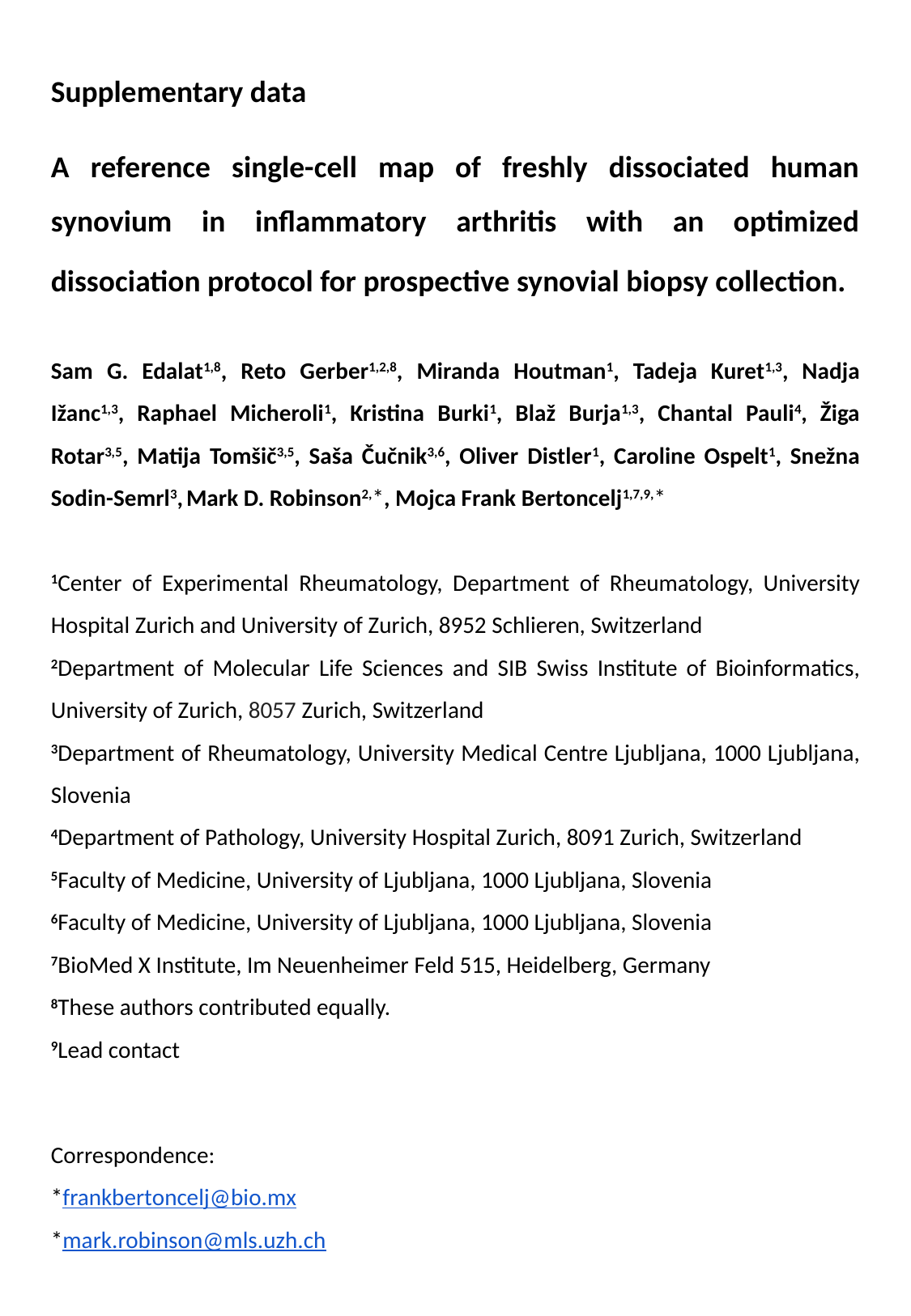

Supplementary data
A reference single-cell map of freshly dissociated human synovium in inflammatory arthritis with an optimized dissociation protocol for prospective synovial biopsy collection.
Sam G. Edalat^1,8^, Reto Gerber^1,2,8^, Miranda Houtman^1^, Tadeja Kuret^1,3^, Nadja Ižanc^1,3^, Raphael Micheroli^1^, Kristina Burki^1^, Blaž Burja^1,3^, Chantal Pauli^4^, Žiga Rotar^3,5^, Matija Tomšič^3,5^, Saša Čučnik^3,6^, Oliver Distler^1^, Caroline Ospelt^1^, Snežna Sodin-Semrl^3^, Mark D. Robinson^2,^*, Mojca Frank Bertoncelj^1,7,9,^*
^1^Center of Experimental Rheumatology, Department of Rheumatology, University Hospital Zurich and University of Zurich, 8952 Schlieren, Switzerland
^2^Department of Molecular Life Sciences and SIB Swiss Institute of Bioinformatics, University of Zurich, 8057 Zurich, Switzerland
^3^Department of Rheumatology, University Medical Centre Ljubljana, 1000 Ljubljana, Slovenia
^4^Department of Pathology, University Hospital Zurich, 8091 Zurich, Switzerland
^5^Faculty of Medicine, University of Ljubljana, 1000 Ljubljana, Slovenia
^6^Faculty of Medicine, University of Ljubljana, 1000 Ljubljana, Slovenia
^7^BioMed X Institute, Im Neuenheimer Feld 515, Heidelberg, Germany
^8^These authors contributed equally.
^9^Lead contact
Correspondence:
*
*

### Slide 2
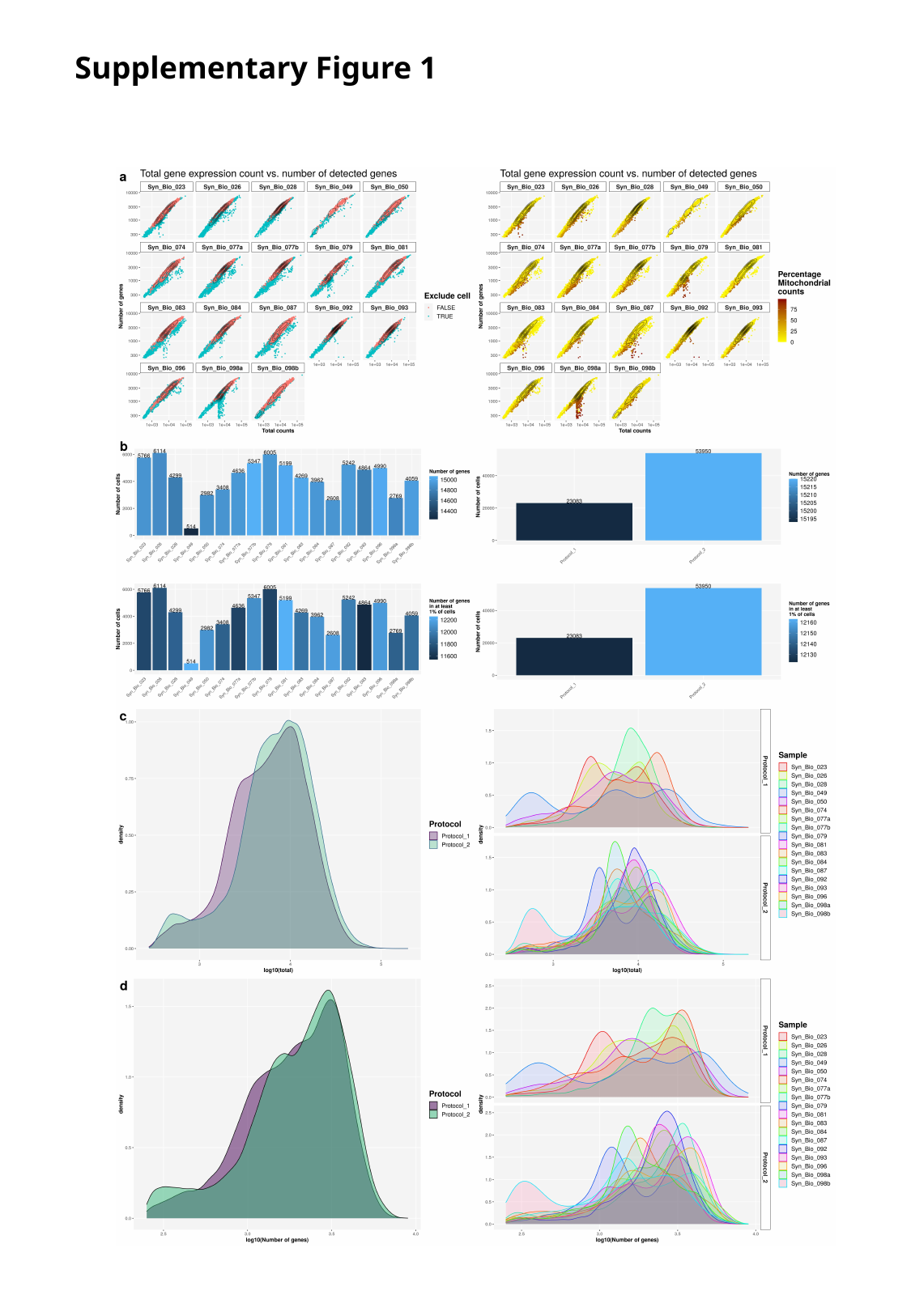

Supplementary Figure 1

### Slide 3
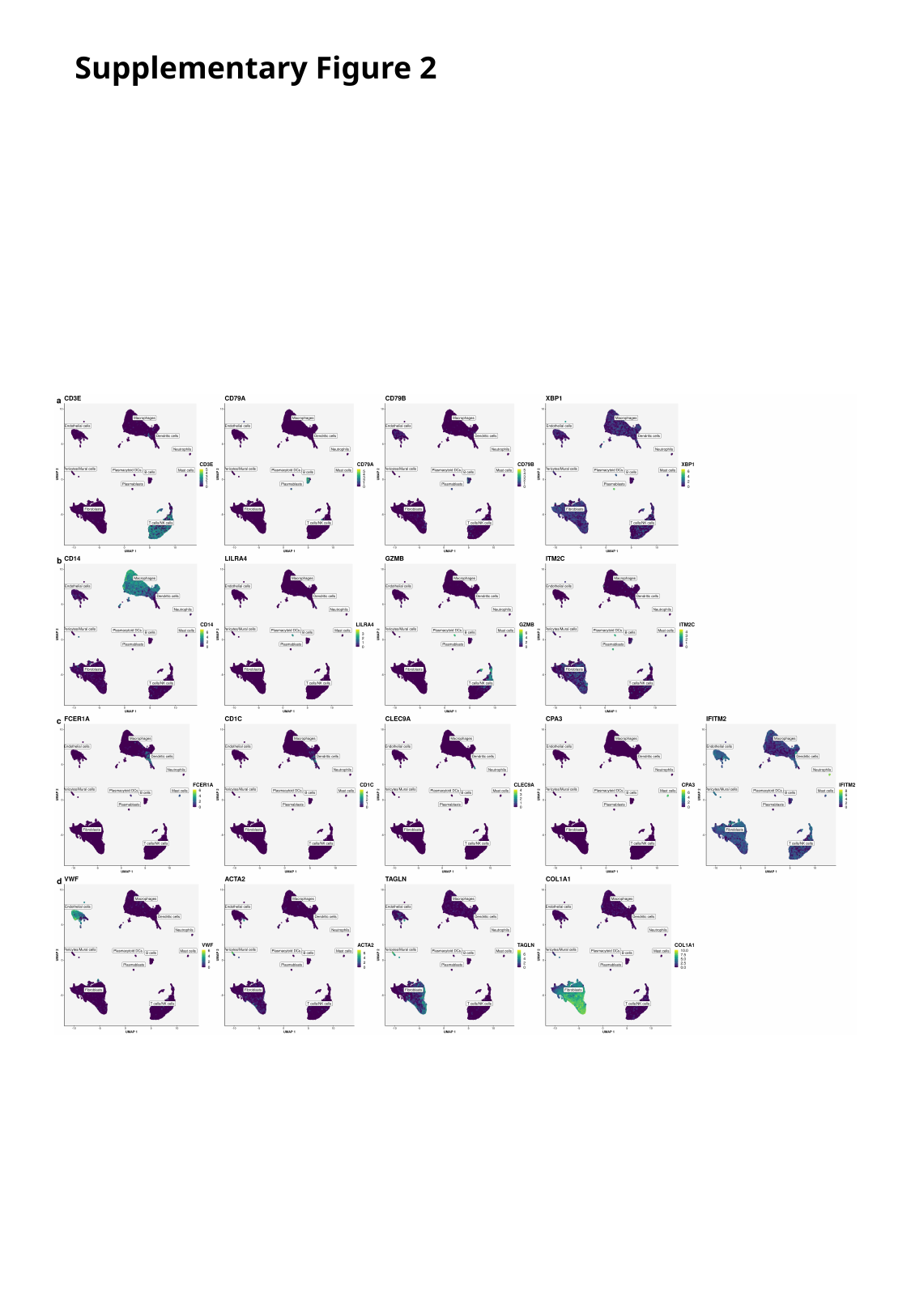

Supplementary Figure 2

### Slide 4
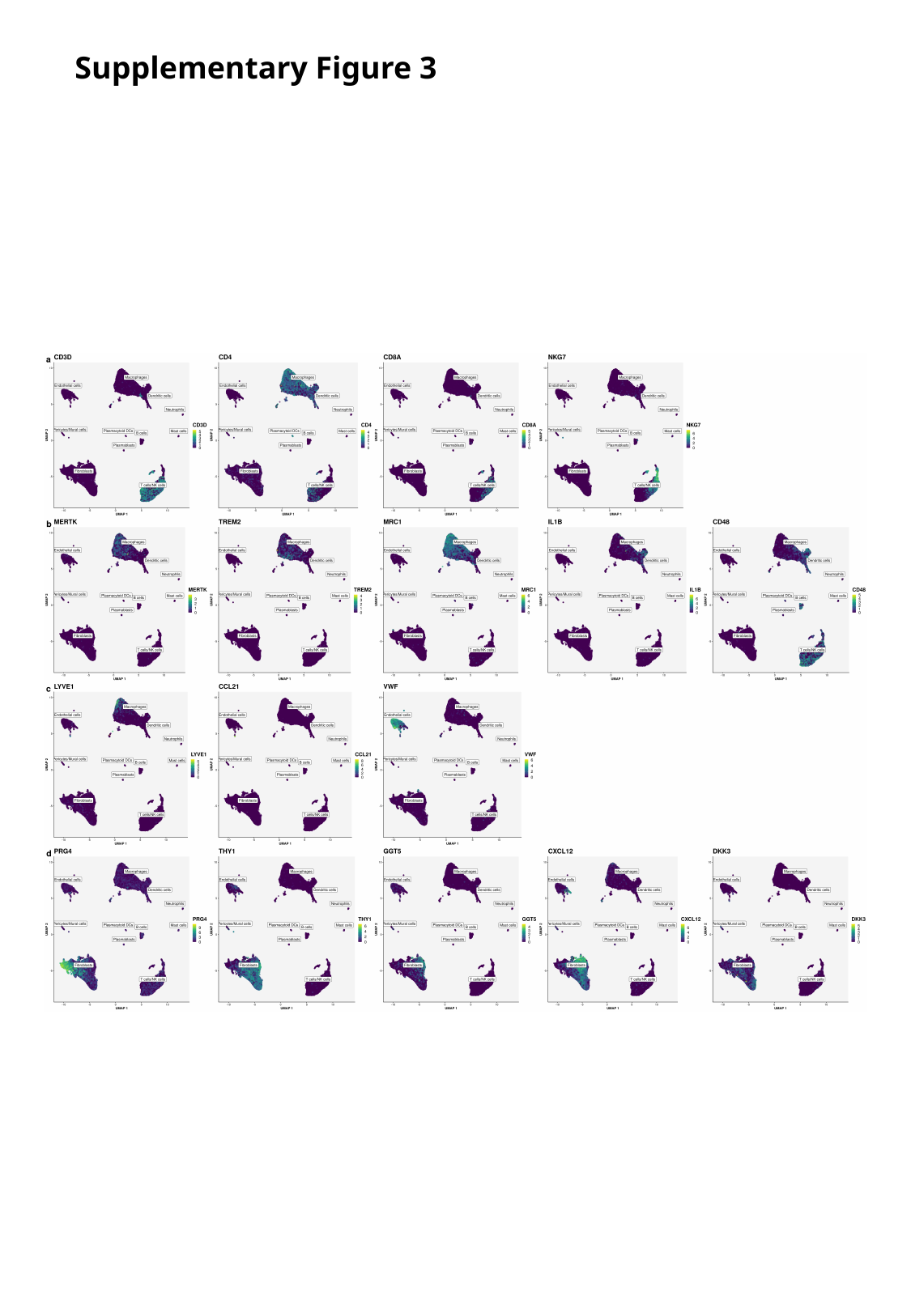

Supplementary Figure 3

### Slide 5
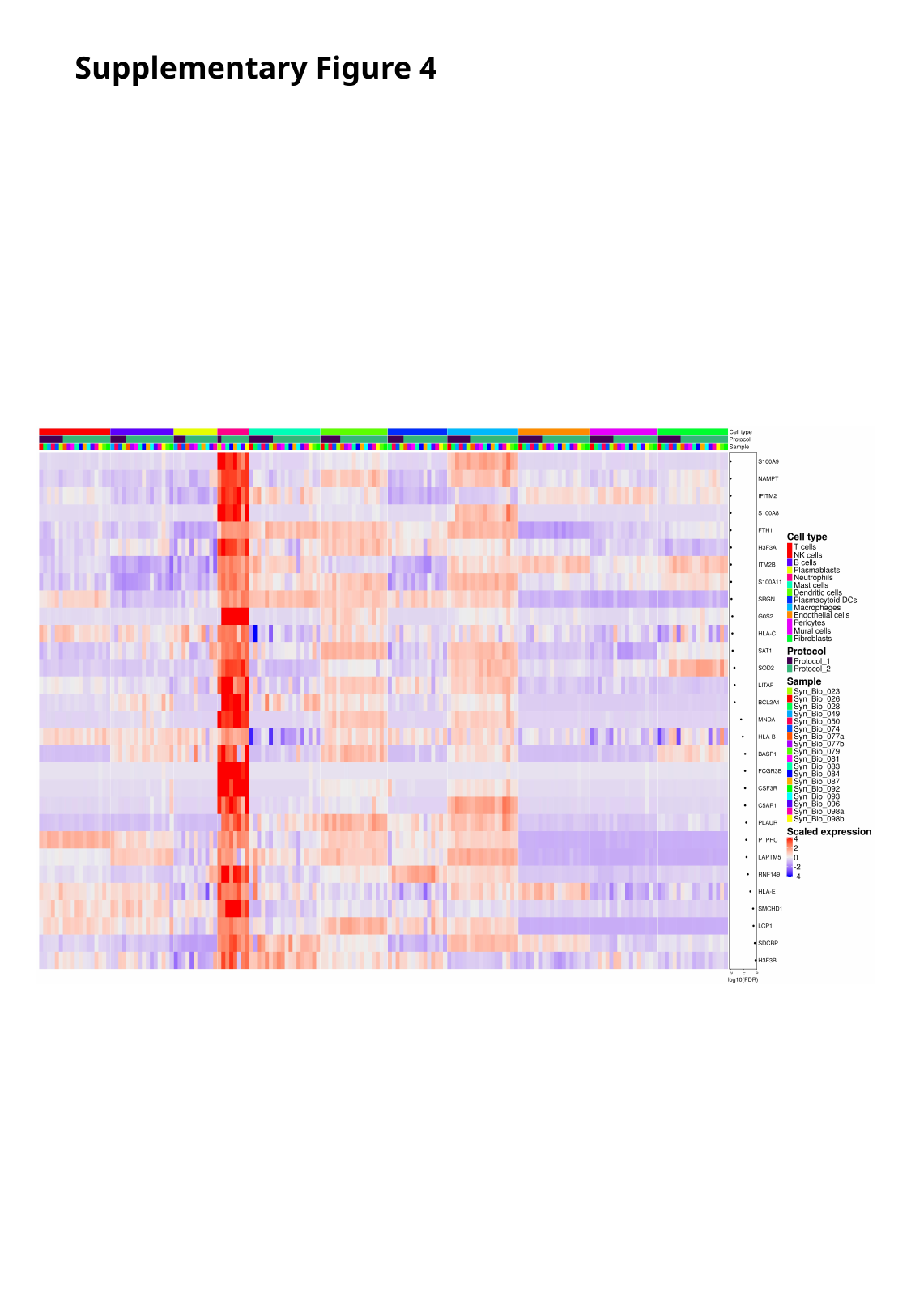

Supplementary Figure 4

### Slide 6
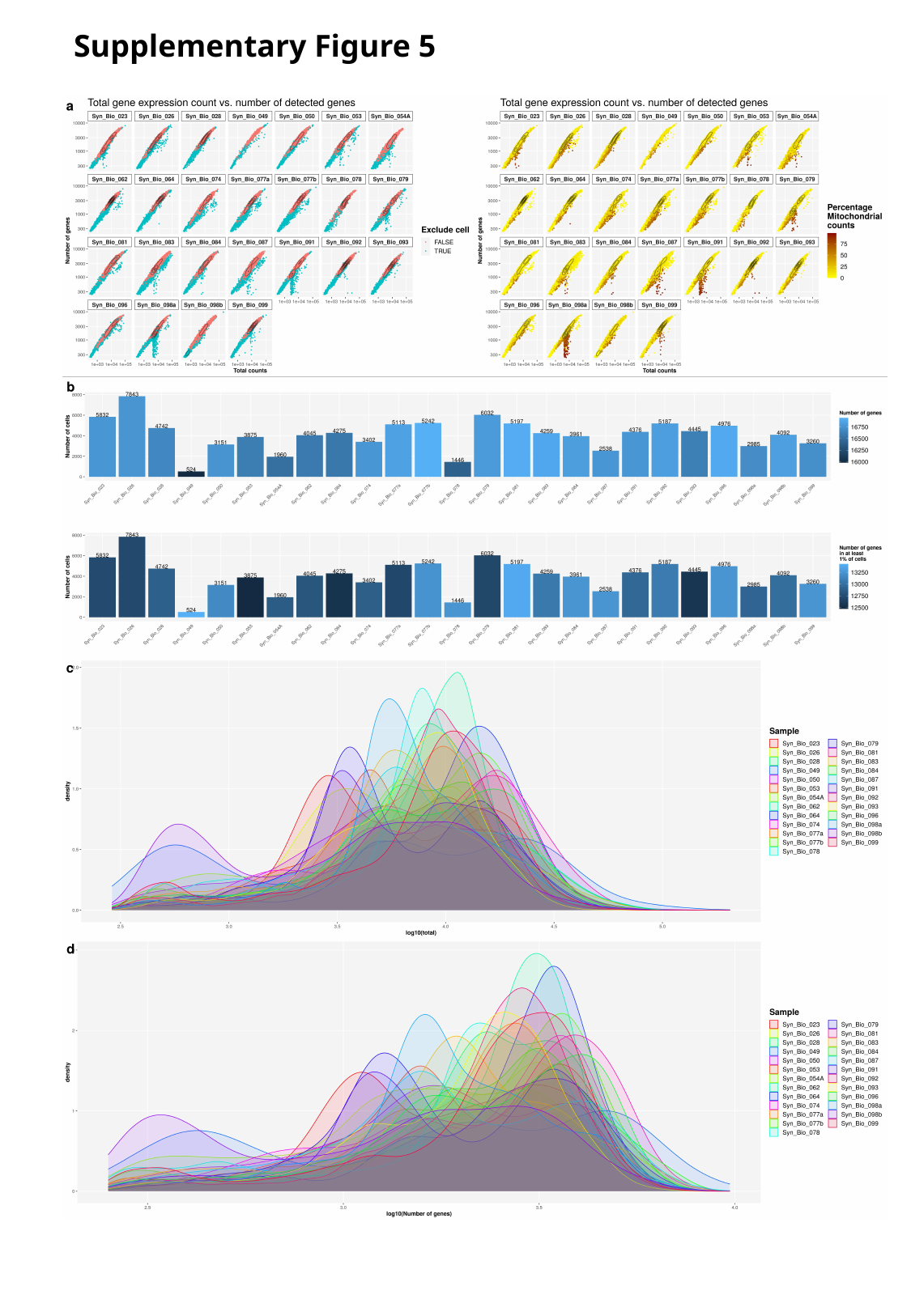

Supplementary Figure 5

### Slide 7
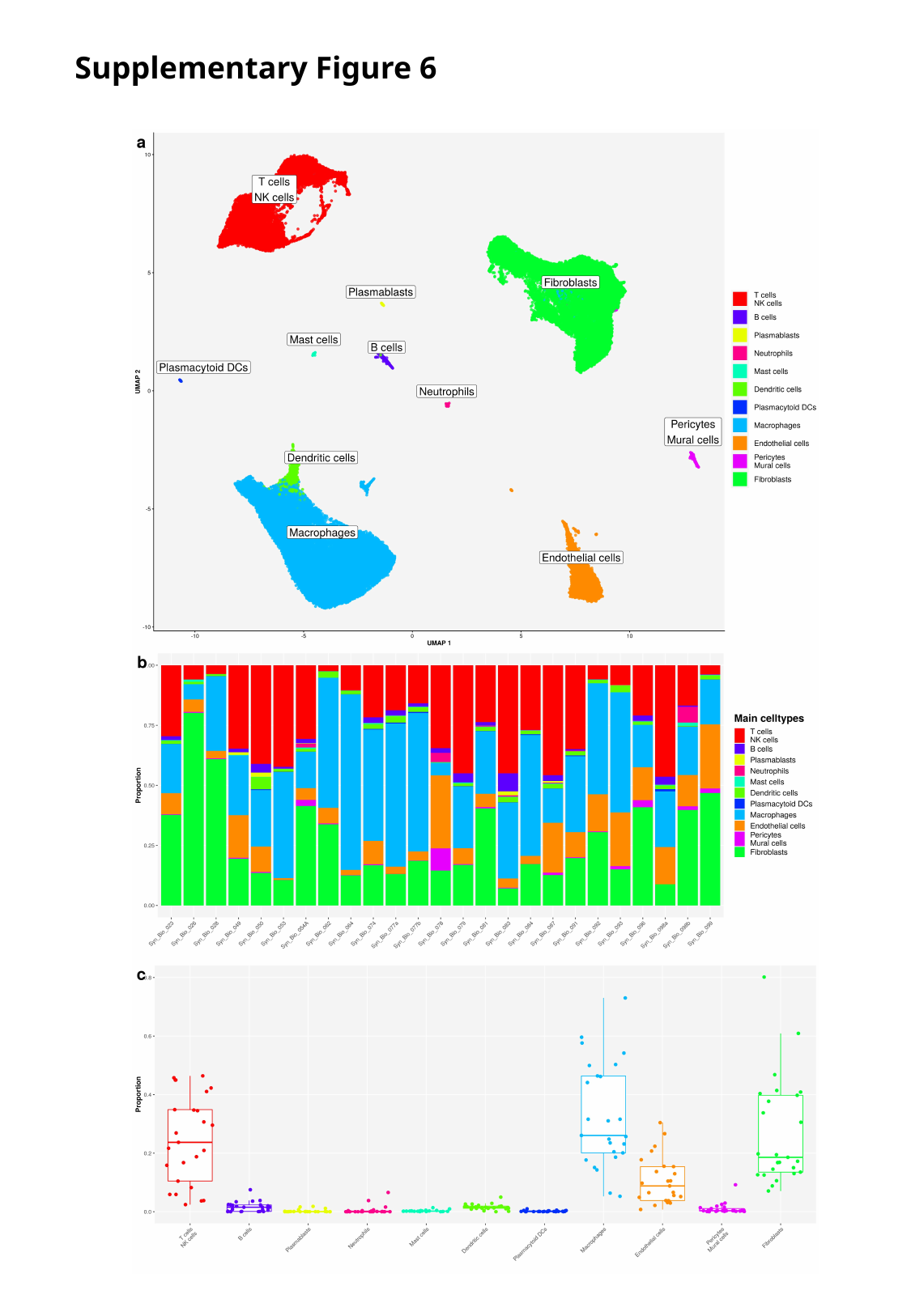

Supplementary Figure 6

### Slide 8
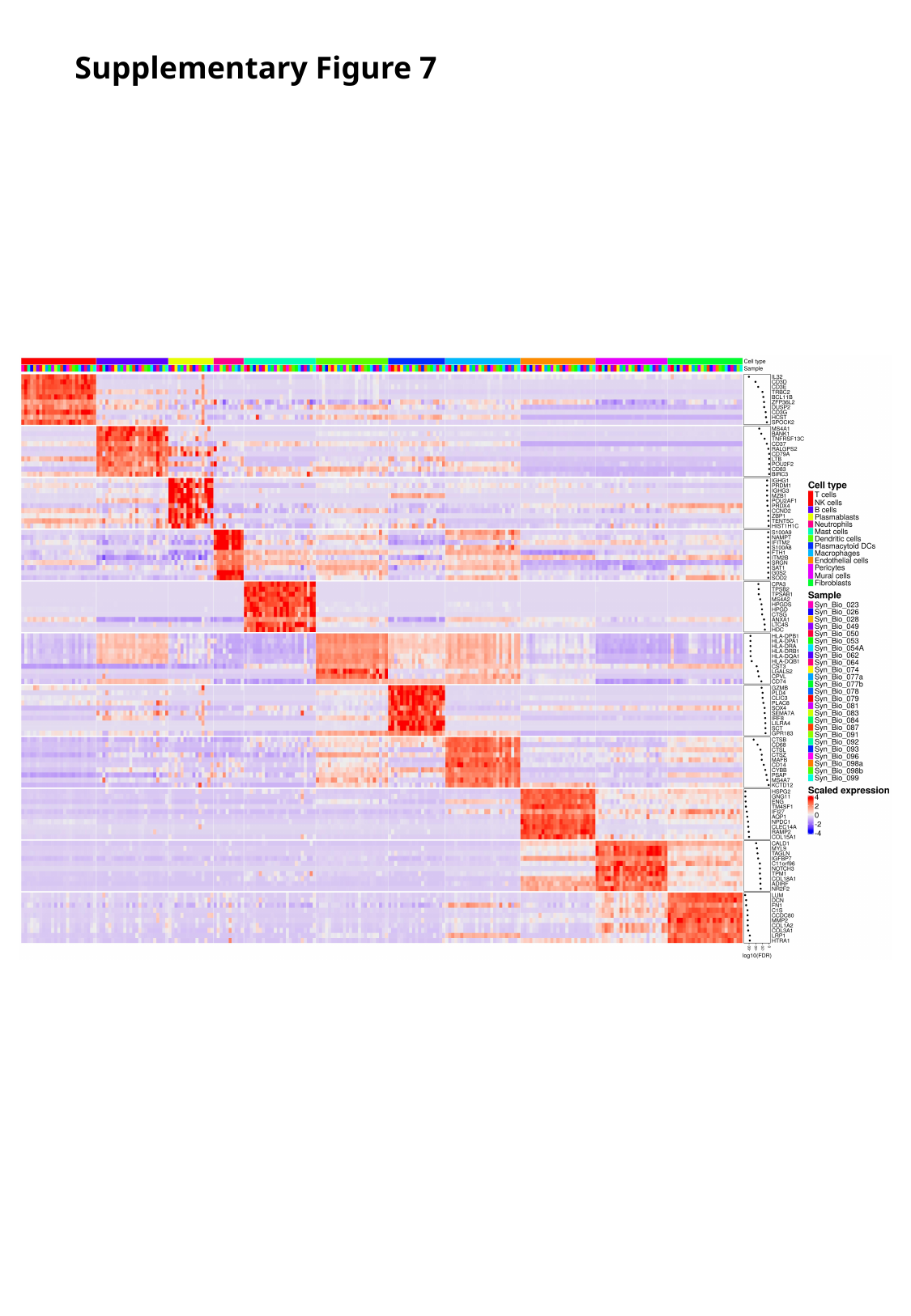

Supplementary Figure 7

### Slide 9
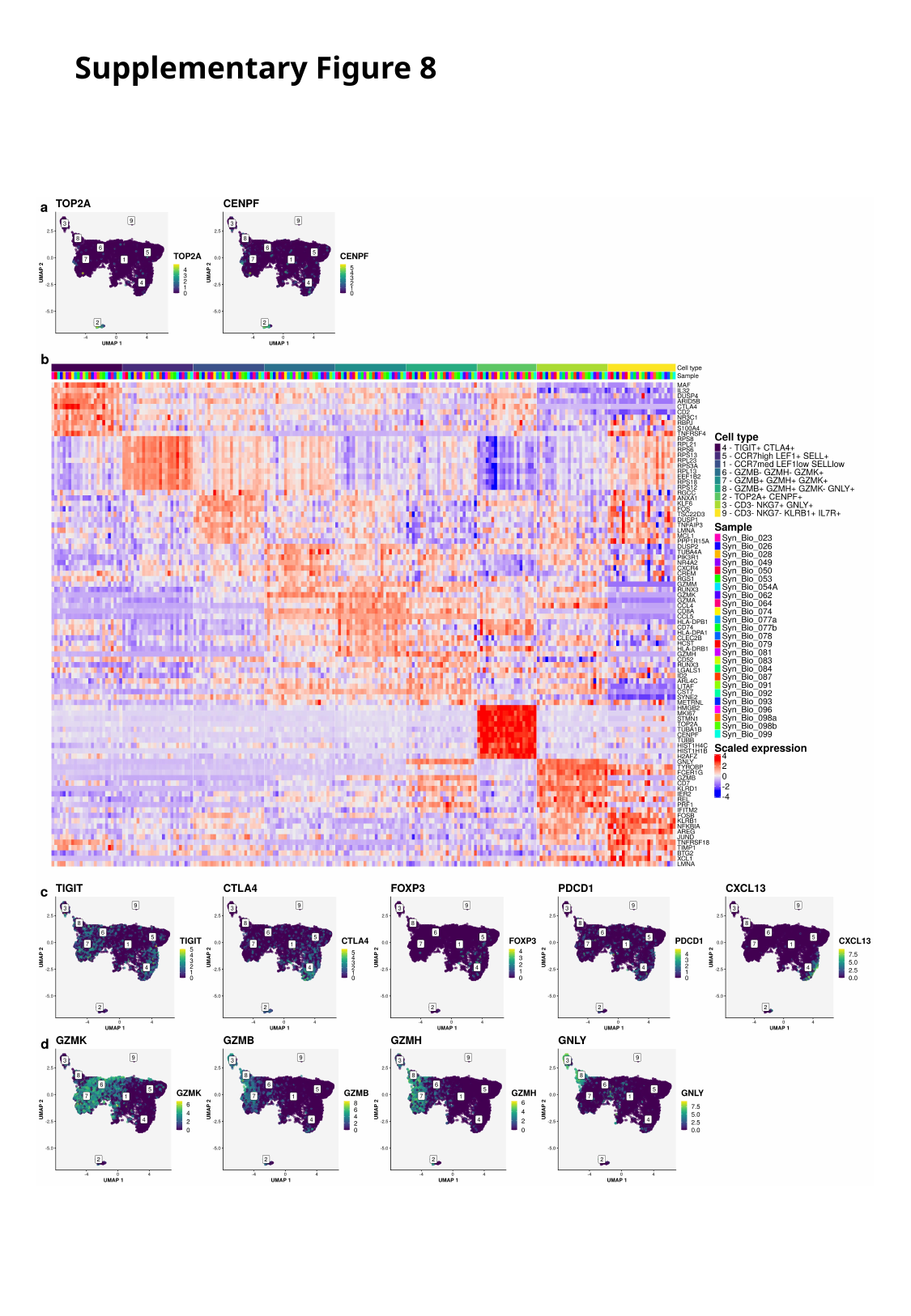

Supplementary Figure 8

### Slide 10
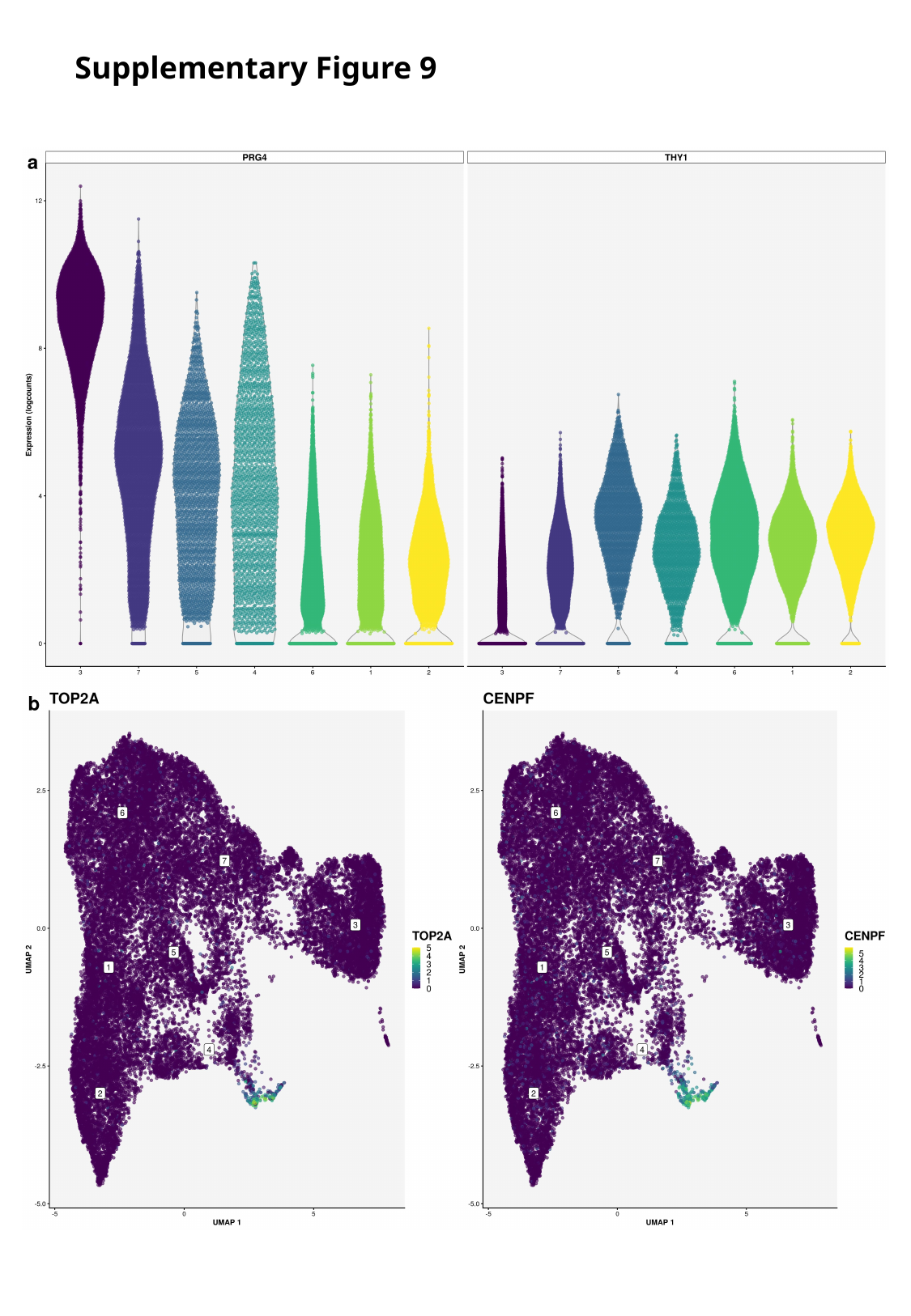

Supplementary Figure 9

### Slide 11
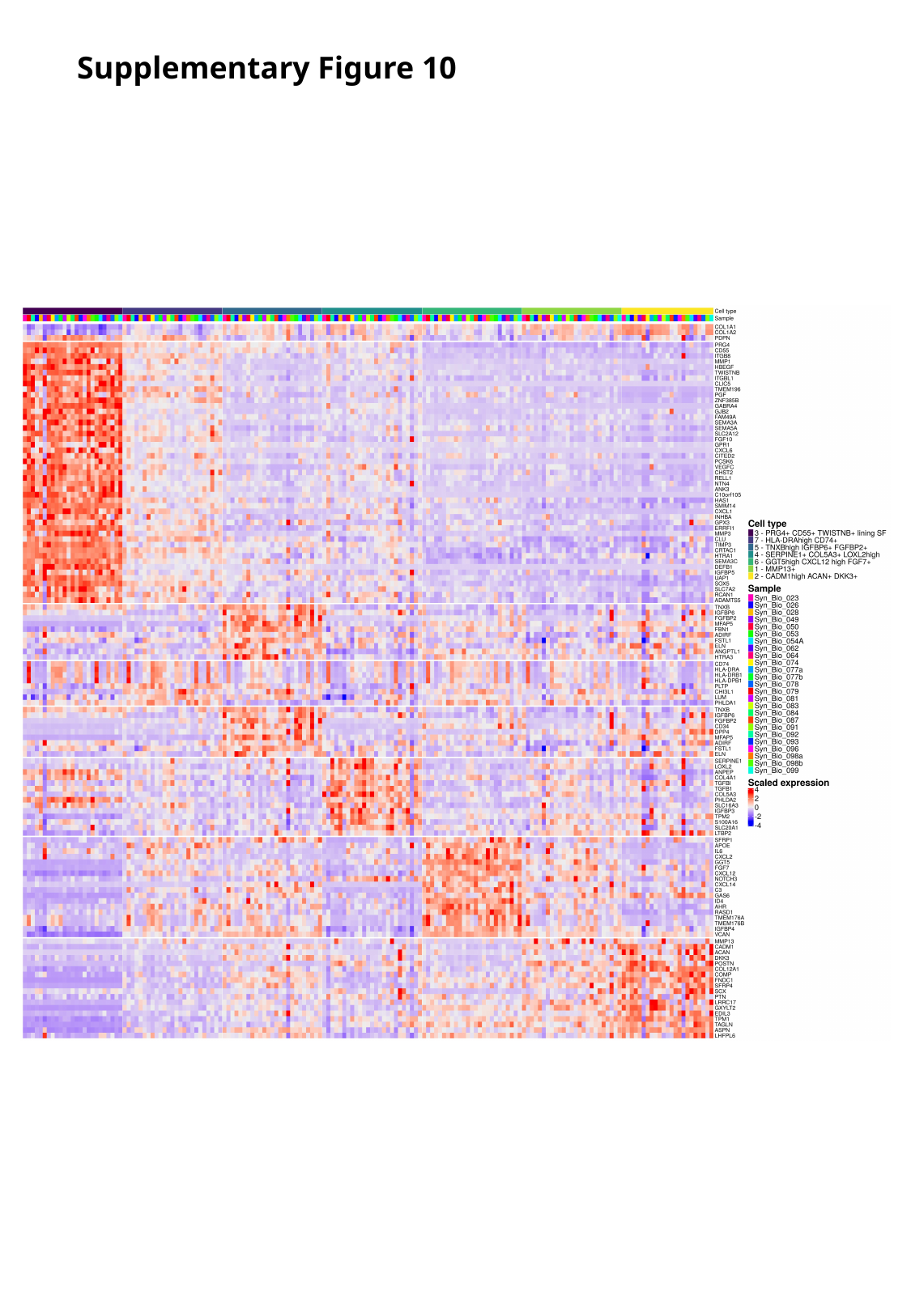

Supplementary Figure 10

### Slide 12
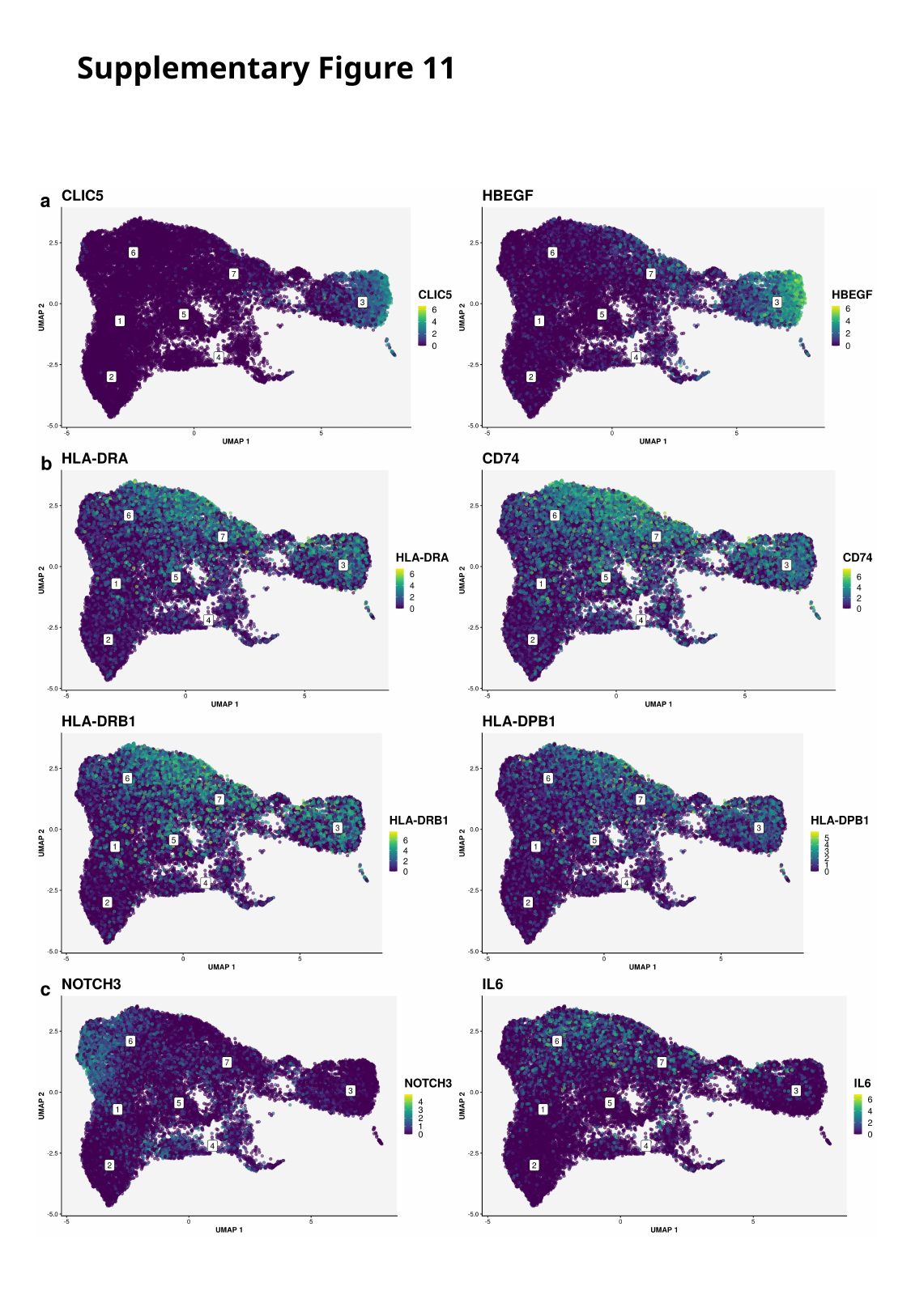

Supplementary Figure 11

### Slide 13
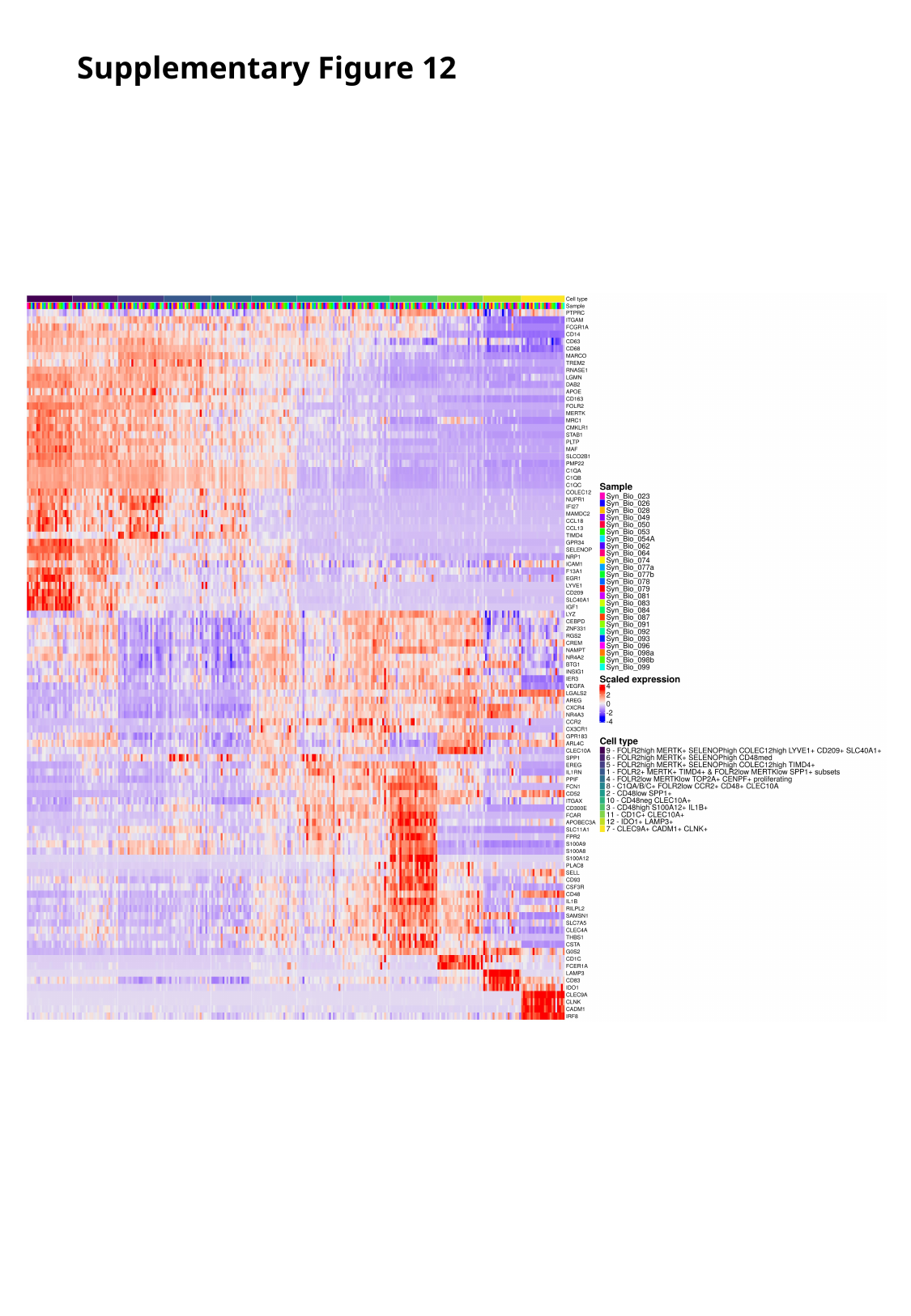

Supplementary Figure 12

### Slide 14
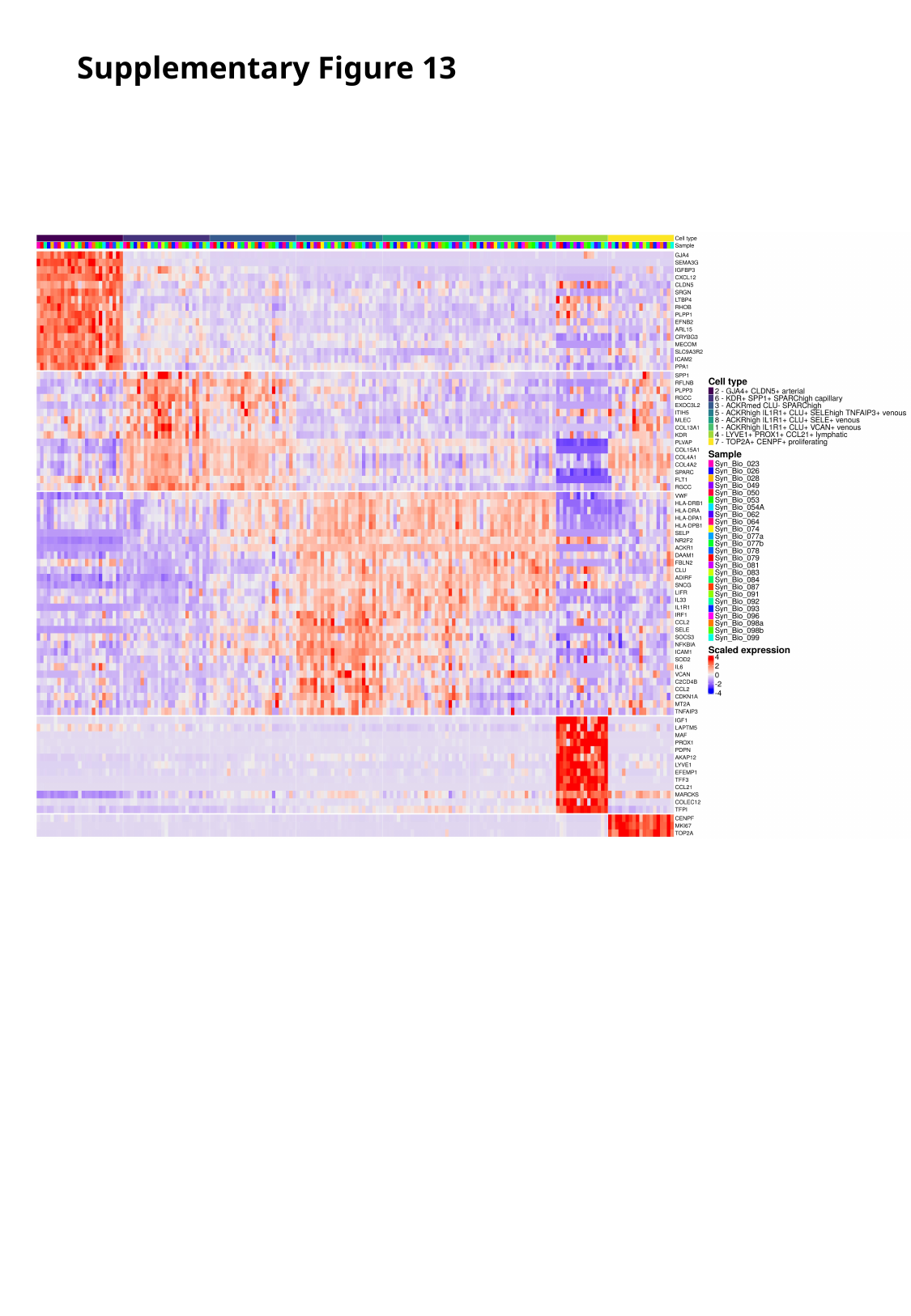

Supplementary Figure 13

### Slide 15
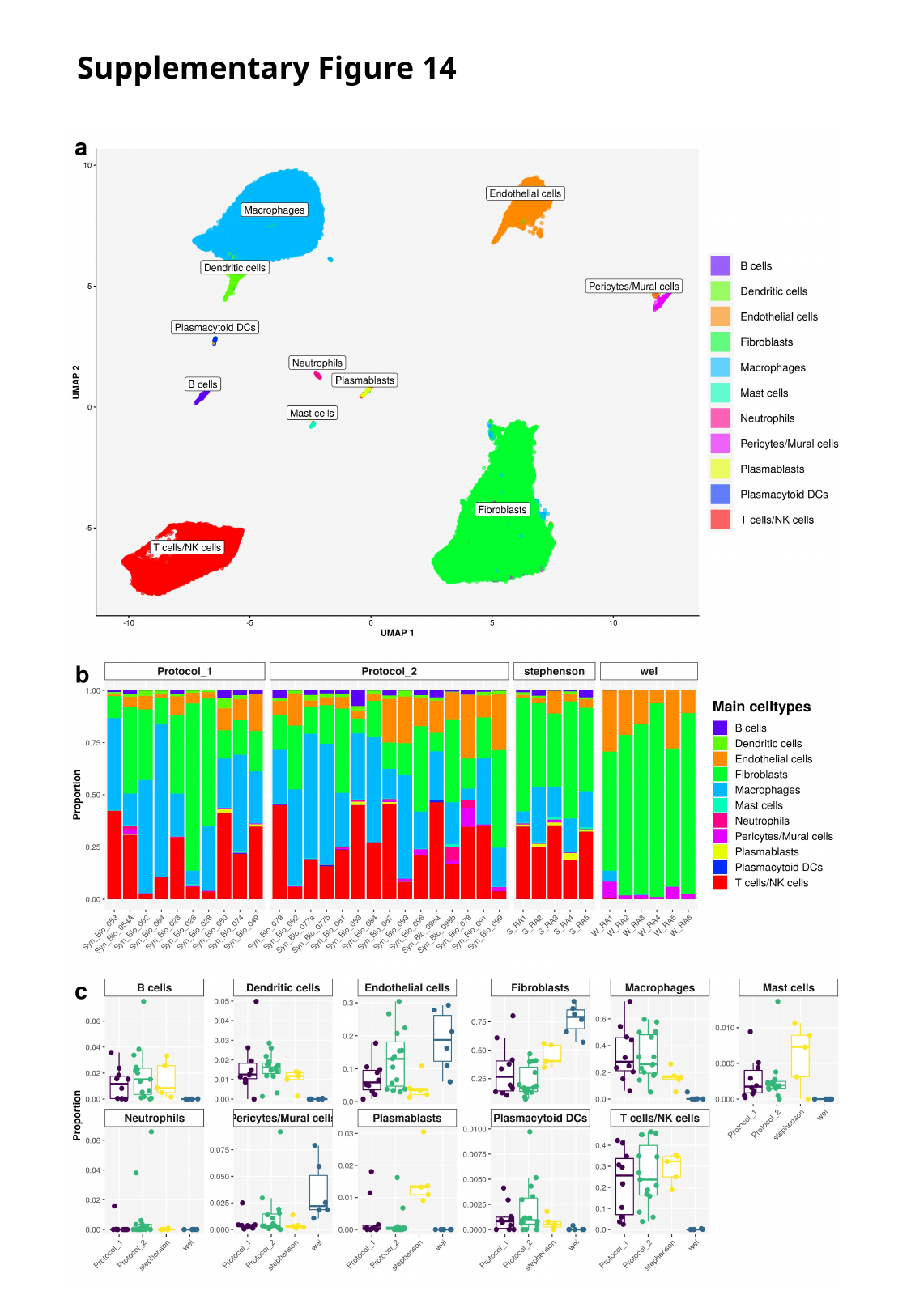

Supplementary Figure 14
